# Cholesteryl Esters Modulate Lipid Droplet Rigidity and Monolayer Organization during Liver Cancer Progression

**DOI:** 10.64898/2026.05.01.722229

**Authors:** Oluwatoyin Campbell, Cecilia Leal, Viviana Monje

## Abstract

In mammalian cells, lipid monolayers support the integrity of lipid droplets (LDs), organelles that function as storage for neutral lipids. Liver-targeting illnesses such as liver cancer interrupt normal LD metabolism and prompt changes in the chemical content of these organelles, which can have effects on structural and organizational behavior of the lipids. In LDs, liver cancer induces concentric crystalline phases of cholesteryl esters (CEs) and triglycerides near the NL-monolayer interface, which become more pronounced as CE concentration increases. Yet, there is little known about how this phenomenon may link to persistence of undigested LDs in liver cancer patients. To shed light on this, all-atom molecular dynamics simulations were used to model LD micropipette aspiration experiments and gain insight into the effect of CE concentration on partitioning, structural, and mechanical properties of LDs. We successfully model micropipette aspiration by application of constant surface tension laterally, which stretches lipid bilayers and monolayers as the magnitude increased. The results show increased phospholipid packing due to insertion of CE fatty tails into the monolayer. Increasing CE concentration induces a non-linear change in surface packing defects on the LDs, notable rigidification, and stiffness. Taken together, these insights improve our understanding of the physical properties at the LD monolayer-core interface during liver cancer progression.

## Introduction

Lipid droplets (LDs) are intracellular fatty bodies which transport energy, stored as triacylglycerol (TAG) and cholesteryl ester (CE) neutral lipids (NLs) (Figure 1A). These lipids are surrounded by a circular phospholipid monolayer that acts as the outer boundary and contains LD surface-associated proteins. The lipid composition and local proteome of these lipid bodies act as modulators of LD structure by influencing rigidity, curvature, and chemical properties which govern LD formation ^1^. LDs play important roles in management of lipid transport and prevention of lipotoxicity, ER stress and autophagy-induced mitochondrial damage, which are important for normal cell function ^2^. LDs are a universal constituent of a variety of tissues such as the muscle, bone, brain, heart and liver. In the liver, LDs play key roles that keep lipid metabolism tightly regulated.

**Figure 1.**
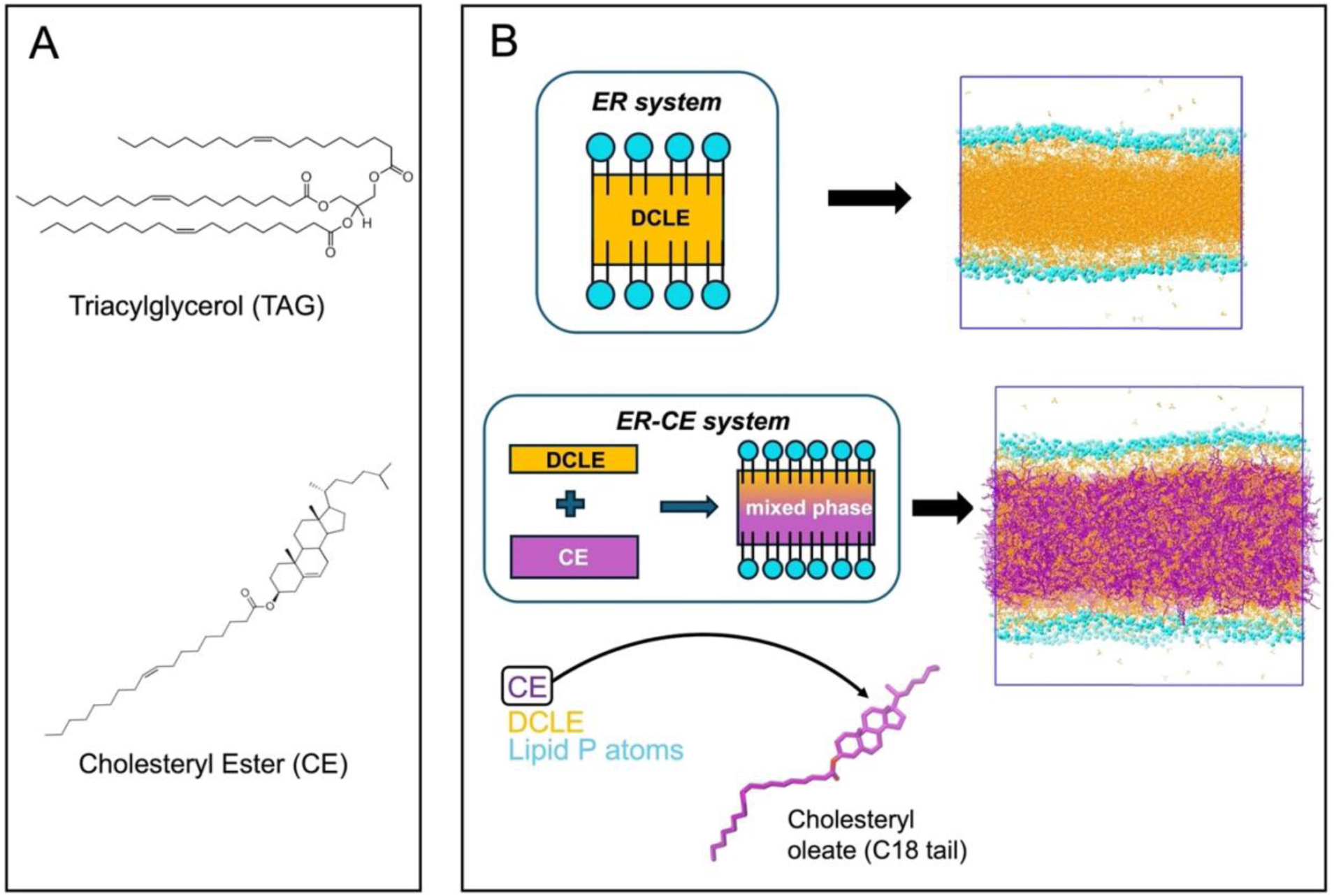
Molecular structure and makeup of lipid droplets (LDs). **A)** Neutral lipids (NLs) found within LDs. Triacylglycerol (TAG) and cholesteryl ester (CE) represented as triolein and cholesteryl oleate respectively. **B)** Illustration of LD model construction. The system contains a trilayer, where phospholipids monolayers (heads in blue) enclose a hydrophobic core containing DCLE hydrophobic solvent (yellow) or a mix of DCLE and cholesteryl oleate (purple). Water and ions hidden for visual clarity.

A common hallmark of liver diseases like viral infection and hepatocellular carcinoma (i.e. primary liver cancer) is the alteration of lipid metabolism, which leads to remodeling of phospholipid and NL content in different ways ^3–8^. These changes tune the cellular environment to favor the spread of these diseases. In liver cancer, for example, acyl-CoA diacylglycerol acyltransferase (DGAT) and acyl coenzyme A cholesterol acyltransferase (ACAT) activity increases TAG and CE content ^9^. In particular, CE abundance has been observed alongside accumulation of lipid droplets and during the late stage of cancers ^9^. Lipidomic and Raman scattering spectroscopy studies have reported increases in CE and proteins involved in its biosynthesis in molecular profiles of hepatocellular carcinoma and other cancer cells ^10–14^.

Experimental and molecular simulations studies have advanced our understanding of LD structure and its ties to NL composition. Recent reports describe how optimal mixtures with mass ratios greater than 4:1 CE:TAG prompt the nucleation of LDs, with lower temperatures promoting a crystalline phase transition at lower CE:TAG compositions ^15, 16^. Cancerous conditions have been shown to drive formation of concentric rings of crystalline CE phases adjacent to the phospholipid boundary ^17^, which may influence lipid transport via alteration of the local lipid landscape where LD surface proteins bind ^18^. Though the monolayer boundary is integral in sustaining the shape of LDs and its interaction with proteins, the implications of LD structure changes are still poorly characterized. To this end, molecular dynamics (MD) simulations, a technique capable of predicting cellular mechanisms involving proteins and lipids based on their structural dynamics and interactions ^19–22^, can be used to probe underlying dynamics and forces that link increased CE content to LD structural integrity and accumulation.

MD simulation studies have focused on elucidating the arrangement and surface properties of lipid molecules in LDs. Increased miscibility of CE with TAG in the NL core and increased presence of TAG within the phospholipid monolayer as CE concentration increases have been reported ^23^. Similarly, TAG sequestration into monolayers prompts packing defects on the LD surface that may aid the binding of necessary LD-associated proteins ^24, 25^. However, previous works have focus on analyzing standard lipid droplet models based on healthy conditions. The specific role of CEs in altering the profile and properties of LD monolayers, and how this may impact LD integrity and longevity due to the onset of liver cancer remains unclear.

To address this knowledge gap, we use all-atom MD simulations to model lipid droplet aspiration experiments with atomistic detail to determine biophysical, structural, and mechanical properties of LDs over the course of progression of liver cancer – characterized by increased CE composition in the bulk phase of the LD core. Using a trilayer model, we probe the effects of monolayer phospholipid composition, CE:NL ratio, and lateral stretching on structural and mechanical properties of the lipid monolayers of constructed LDs. This generated over 40 microseconds of data, which shows that increase in CE content promotes a rigidification of LDs as interdigitation of the molecules increasingly manifests. The structure of the monolayer becomes more ordered, with increased CE presence in the monolayer driving water repulsion from the monolayer-water interface. These observed changes have implications in the stiffness and targeting of LD proteins to these organelles, making our study useful for the development of targeted cancer therapeutics.

## Methods

Increase in CE composition is a key indicator of advancement in liver cancer ^9, 10, 12^. Here, we evaluate the effects of cholesteryl oleate on the partitioning, biophysical and mechanical properties of LD monolayers. To accomplish this, 4 different models were designed and studied with MD simulations: (i) the *Pure* model containing DOPC monolayers and the hydrophobic solvent, 1,1-dichloroethane (DCLE), to represent the NL phase of LDs; (ii) the *ER* model containing monolayers made up of DOPC, DPPE and POPI in a 62:24:14 mol% ratio and a DCLE NL phase; (iii) the *low-CE* model containing the *ER* monolayer and a 60:40 CE:DCLE mass% (18:82 mol%) mix within the hydrophobic NL phase; and (iv) the *high-CE* model containing the *ER* monolayer and a 85:15 CE:DCLE mass% (46:54 mol%) mix within the hydrophobic phase. DCLE has been used to imitate the membrane hydrophobic interior and increase diffusion of adjacent molecules to speed up reorganization processes required for biomolecular mechanisms of membrane proteins ^26^. This made it an appropriate solvent to accelerate diffusion dynamics of CE molecules; additional information about the systems in this study is summarized in Table S2.

The LD models were constructed as trilayers, similarly to models explored by other groups ^24, 25, 27, 28^. These layers include two individual leaflets which sandwich the hydrophobic phase that takes the form of a simulation box populated by desired representative molecular mixtures (Fig. 1B). First, leaflets were built and equilibrated in bilayers containing 400 lipids per leaflet for the *Pure* and *ER* systems, and 600 lipids for the *low-CE* and *high-CE* systems using CHARMM-GUI Membrane Builder ^29, 30^ and GROMACS ^31^. The DCLE *gro* structure file was attained from CHARMM-GUI HMMM Builder ^26^, and the solvent boxes built using GROMACS. The structure of cholesteryl oleate was generated with Avogadro ^32^ and parametrized with the CGenFF tool ^33^ on CHARMM-GUI Solution Builder ^34, 35^. The layers were assembled with a Bash script and GROMACS built-in commands; modeling scripts, sample files, and associated directions can be found at https://github.com/monjegroup/LD-structure.

Simulations of the models comprised of the bilayer and NL box equilibration, the trilayer NPT production, and the trilayer N𝑃_𝑧_γT production, where γ represents a constant surface tension applied laterally to stretch the lipid layer plane. For each replica, the bilayer equilibration was initiated with the 6-step equilibration protocol generated by the Membrane Builder upon system build. This involves a steepest descent minimization to less than 1000 kJ mol^-1^ nm^-1^, two steps of 125 ps NVT, one step of 125 ps NPT and finally 3 steps of 500 ps NPT, all accompanied with gradual reduction of applied lipid position and dihedral restraints. Finally, the 200 ns equilibration was run.

The NL simulation boxes equilibration involved a 1000 kJ mol^-1^ nm^-1^ steepest descent minimization, one step of 100 ps NVT and one step of 100 ps NPT. Using the appropriate equilibrated bilayer and NL box coordinates, each trilayer system model was assembled by stacking the bilayer bottom leaflet, the NL box and the bilayer top leaflet within a new simulation box, then populating the vacuum space with water and neutralizing ions using 0.15 M KCl. Water and ions placed between the leaflets (i.e. in the hydrophobic phase) were manually deleted. The trilayer NPT production phase was carried out for 100 ns on the *Pure* and *ER* model systems, 500 ns on the *low-CE,* and 1 μs on the *high-CE* model systems. In some cases, a short NPT simulation of 5-80 ns was run before this step to further stabilize the beginning of the production phase.

The N𝑃_𝑧_γT production phase was run for 200 ns, 500 ns, 1 μs and 2 μs on the *Pure*, *ER*, *low-CE* and *high-CE* models, respectively, and a constant surface tension value between 2-32 mN/m (see Table 1 for tension settings of each model). A minimum tension of 0.625 mN/m was added to these values to prevent structure collapse, based on the non-zero surface tension measurements in the bilayer model simulations. Altogether, a total of 44.4 μs of data were generated, providing ample sampling for statistical analysis and evaluation of structural and mechanical properties induced by lateral stretching as well as changes in lipid composition. Detailed settings and conditions for each simulation phase are summarized in the first section of the supporting information (SI).

**Table 1.**
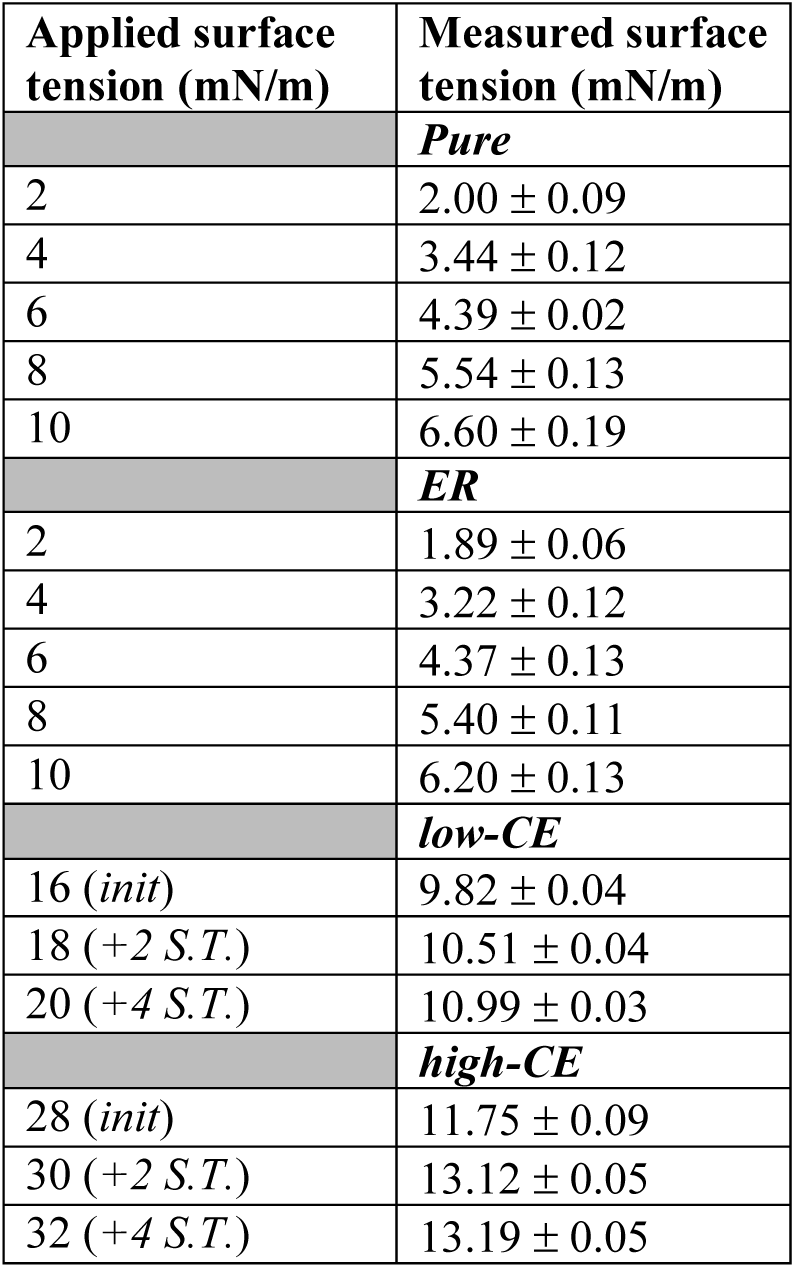
Measured surface tension in LD models. The applied surface tension is the value entered into the simulation run parameter file, and the measured surface tension is the one calculated from the completed simulation.

Trajectories visualization and snapshot rendering were performed with VMD ^36^. Data analysis and plotting were carried out with GROMACS, VMD, and Python (Pandas, Numpy, Seaborn, and MDAnalysis ^37, 38^ libraries). When delta (Δ) is present in the name of the property plotted, the reported values are the difference calculated between the measured value for the stretched case and that of a bilayer with similar composition (*Pure* and *ER*) or the unstretched trilayer (*low-CE* and *high-CE*). Description of key analyses featured in this work can be found in second section of the SI.

## Results and Discussion

### Modeling lipid droplet aspiration

Micropipette aspiration is a well-established method for determining mechanical and viscoelastic properties of cells and other lipid-enclosed materials ^39, 40^. It involves application of a suction pressure to examine the deformation response of an interface. A pseudo-form of this approach is applied in this work by gradually increasing the surface tension along the X and Y directions of the trilayers, thereby inducing stretching along the monolayer plane.

To determine the optimal size for the monolayer leaflets, bilayers with 200 and 400 lipids/leaflet (L/L) were equilibrated for 200 ns and used to construct the trilayer models. Figure S1 shows the time series of the measured surface tension when set at 2, 4, 6, 8 and 10 mN/m. At low surface tensons of 2 and 4 mN/m, the trilayer systems buckle under 200 L/L conditions but remain stable in 400 L/L systems (Fig. S1A). Buckling prompted eventual collapse due to a sudden horizontal compression and elongation in the Z dimension of the simulation box, compared to stable simulations characterized by horizontal stretch and vertical compression (Fig. S1B). In a multiscale MD simulation study, decrease in monolayer leaflet size reduced long-wavelength undulations and increased repulsion as indicated with a lateral pressure profile analysis ^24^. It is likely that this disruptive phenomenon is at play within the 200 L/L models, acting to prevent favorable coexistence of the interfaces. The measured surface tension from the equilibrated trajectories was lower than the set value, albeit by the same amount with increasing surface tension (Table 1).

Since larger leaflets prevented collapse of the LD models, we designed CE-containing models to include 600 L/L. Running at surface tensions lower than 16 mN/m for *low-CE* models and 28 mN/m for *high-CE* models lead to similar collapse in Fig. S1B. Therefore, incremental tension was applied to these base values, referred to as the *init* case in Table 1 for simplicity. The 18 and 20 mN/m cases in the *low-CE* model, and 30 and 32 mN/m cases in the *high-CE* model are referred to as *+2 S.T.* and *+4 S.T.* cases respectively. From here on, these names and applied tensions are used to describe results.

### Structure modulation and molecular partitioning in the absence of CE

The mechanical response of cell organelles depends on the ratio of lipid species and their interactions with one another ^41, 42^. The physical response and integrity of LDs due to phospholipid composition were evaluated using various structural analyses shown in Figure 2. The area per lipid (APL) in *Pure* and *ER* models increase with tension (Fig. 2A). The splay angle of phospholipids in both models also increases in a similar fashion (Fig. 2B-C). Deuterium order parameters (S_CD_) were measured to evaluate packing of lipid tails. To aid interpretation of the behavior of acyl tails of both phospholipids and CE, the values plotted are multiplied by -1 (equation 4 in SI). -S_CD_ decreases for each carbon in both models, showing more disorder (i.e. less well-packed conditions) as stretching occurs (Fig. 2D-E, Fig. S2). Interestingly, the difference in splay angle and tail packing between the *ER* monolayer and its bilayer counterpart is larger compared to the *Pure* model. These results suggest both stretching and phospholipid composition have a non-trivial impact on the structural conformation and lateral packing of lipid tails.

**Figure 2.**
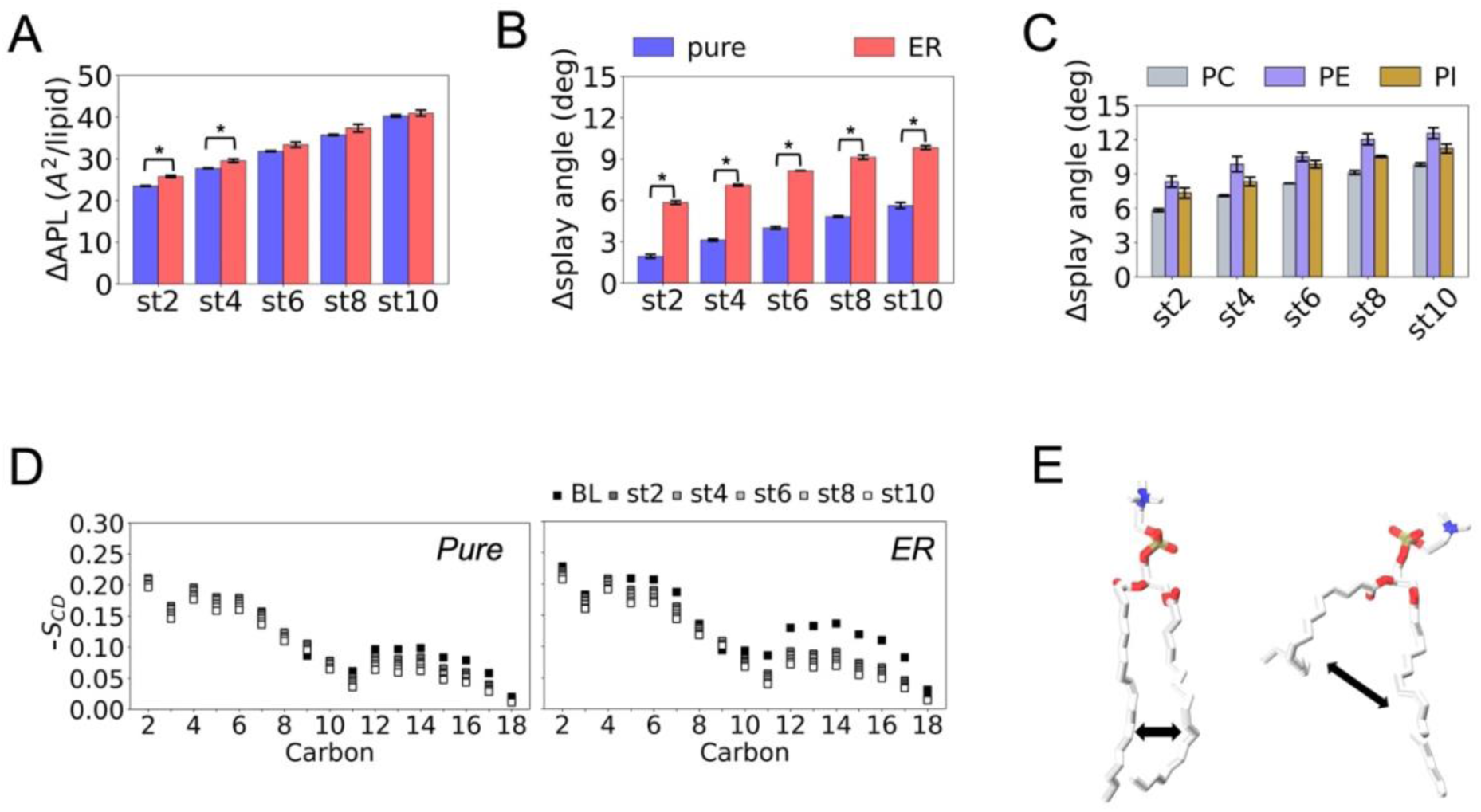
Structure of monolayer lipids in the absence of cholesteryl ester (CE). **A)** Area per lipid. **B)** PC lipid splay angle. **C)** Comparison of PC, PE and PI lipid splay angles. **D)** Deuterium order parameters (S_CD_) of PC lipid sn1 tails. **E)** Representative snapshot of randomly selected PC lipid showing increased disorder, which involves larger splay and bending in acyl tails. Larger -S_CD_ indicates more ordered packing, smaller -S_CD_ indicates more disordered packing. Delta (Δ) indicates the difference calculated between the measured value and that of a bilayer with similar lipid composition. Standard error across replicas indicated with error bars, and “*” indicates significant difference in means (p < 0.05) between compared models (*pure* and *ER*).

Next, the electron density profile (EDP) of water and DCLE were calculated to examine the partitioning of non-lipid constituents based on distribution of electrons within each molecule. As tension increases, slight thinning occurs in the *Pure* (Fig. S3A-B) and *ER* EDP distribution (Fig. 3A-B). This manifests as both vertical compression and stretching (Fig. S4).To quantify the effect of compression, lateral pressure profiles (LPPs) were calculated using the GROMACS-LS software ^43^. Results corroborate the thinning behavior of the EDP, indicated the inward shift of the curves as tension increases (Fig. 3C). For all models, conserved positive peaks in the LLPs represent repulsive forces at the water-headgroup and tail-DCLE interfaces, while the negative minima represent attractive forces within the lipid tail region of the monolayer. Minor peaks appear in between the largest peaks and the center of the systems, indicating increased repulsion at the phospholipid-DCLE interface. Water permeation was also probed by measuring the difference between the phospholipid headgroup Z position (entire system EDP peak) and the Z position where the water EDP becomes effectively zero. The permeation decreases with increasing tension, though no statistical difference is seen between the *Pure* and *ER* models at the same applied tension (Fig. 3D).

**Figure 3.**
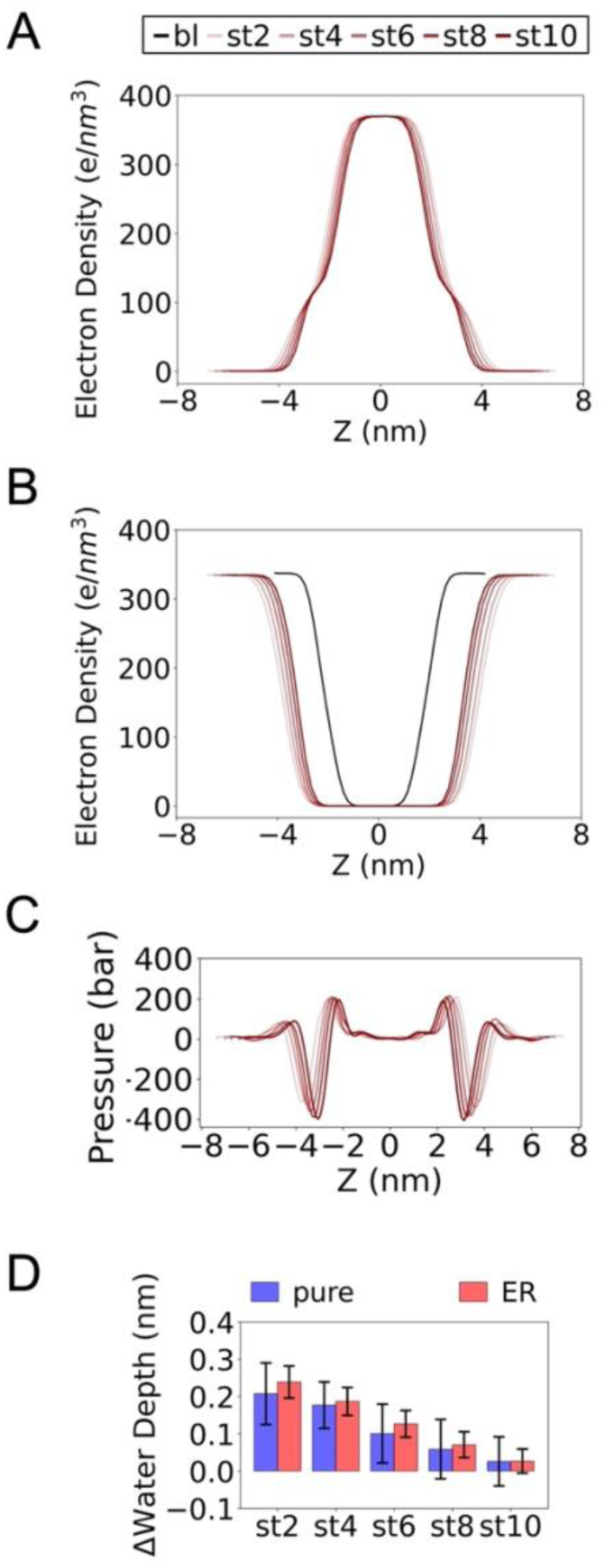
Distribution and dynamics of molecular components in the absence of CE. **A)** Electron density profile (EDP) illustrates partitioning behavior of DCLE and **B)** water molecules in the *ER* model. **C)** Lateral pressure profile (LPP) in the *ER* trilayer model. **D)** Water depth past phosphate heads in the *Pure* and *ER* monolayers. Results of increasing tension symbolized by a gradient of light to dark red. Standard error across replicas indicated with error bars. No significant difference in means (p < 0.05) between compared models (*pure* and *ER*).

These results show that repulsion between solvent and phospholipid molecules can build up due to stretching and vertical thinning at the monolayer surface. Both *Pure* and *ER* models experience near exact trends in partitioning, suggesting a low impact of phospholipid composition on these properties.

### Determination of convergence in CE-containing models

So far, noticeable differences in lipid tail structure are seen upon changes in phospholipid composition, while little effect is observed in component partitioning. In subsequent sections, we adopt the more complex ER model to elucidate the effects of CE on the mechanical properties and organizational behavior at the hydrophobic region of the LD monolayer.

To determine lateral dynamics within the models, we evaluated the lipid diffusion coefficient as estimated from the mean square displacement of each lipid. Diffusion of PC lipids increases as stretching becomes more pronounced in the *Pure* and *ER* models, while it remains unchanged in *low-CE* and *high-CE* models (Fig. S5A); this trend is captured for all lipid types (Fig. S5B). Concurrently, increase in CE composition induces a decrease in diffusion. This implies that the presence of CE interferes with the increased spatial freedom (i.e. APL) induced by stretching (Fig. S5C).

Equilibration was defined when the APL time series of the monolayers becomes constant. This takes less than 20 ns for *Pure* and *ER* models, 50 ns for all *low-CE* runs, 700 ns for *high-CE* unstretched (*npt*) runs, and at least 1,250 ns for *high-CE* stretched runs (Fig. S5C). Despite the reduction in lipid diffusion due to CE, the densest *high-CE* model equilibrates within 1.25 μs out of a total of 2 μs, ensuring over 700ns of trajectory for analysis.

### Structure modulation and molecular partitioning in the presence of CE

Using the same structural analyses featured in Figure 2, we examined the CE-containing models. Figure 4A shows a marked increase in the APL with increasing tension and CE abundance. The splay angle also increases with CE concentration (Fig. 4B-C). Interestingly, stretching fosters increase in the splay angle for the *low-CE* model, but has no effect in the *high-CE* model. Figure 4D shows decrease in the -S_CD_ of PC lipids upon stretching. The values in *high-CE* are slightly less than those of the *low-CE* model at the same applied tension for C2-C10, but higher than *low-CE* for C11-17 in the sn1 tails. Similarly to trends in splay angle, the increasing tension in *high-CE* induces little to no change in the order parameters (Fig. 4D, right panel). These trends are sustained for the other lipids (Figs. S6 and S7), showing that high CE abundance stifles lipid conformational changes.

**Figure 4.**
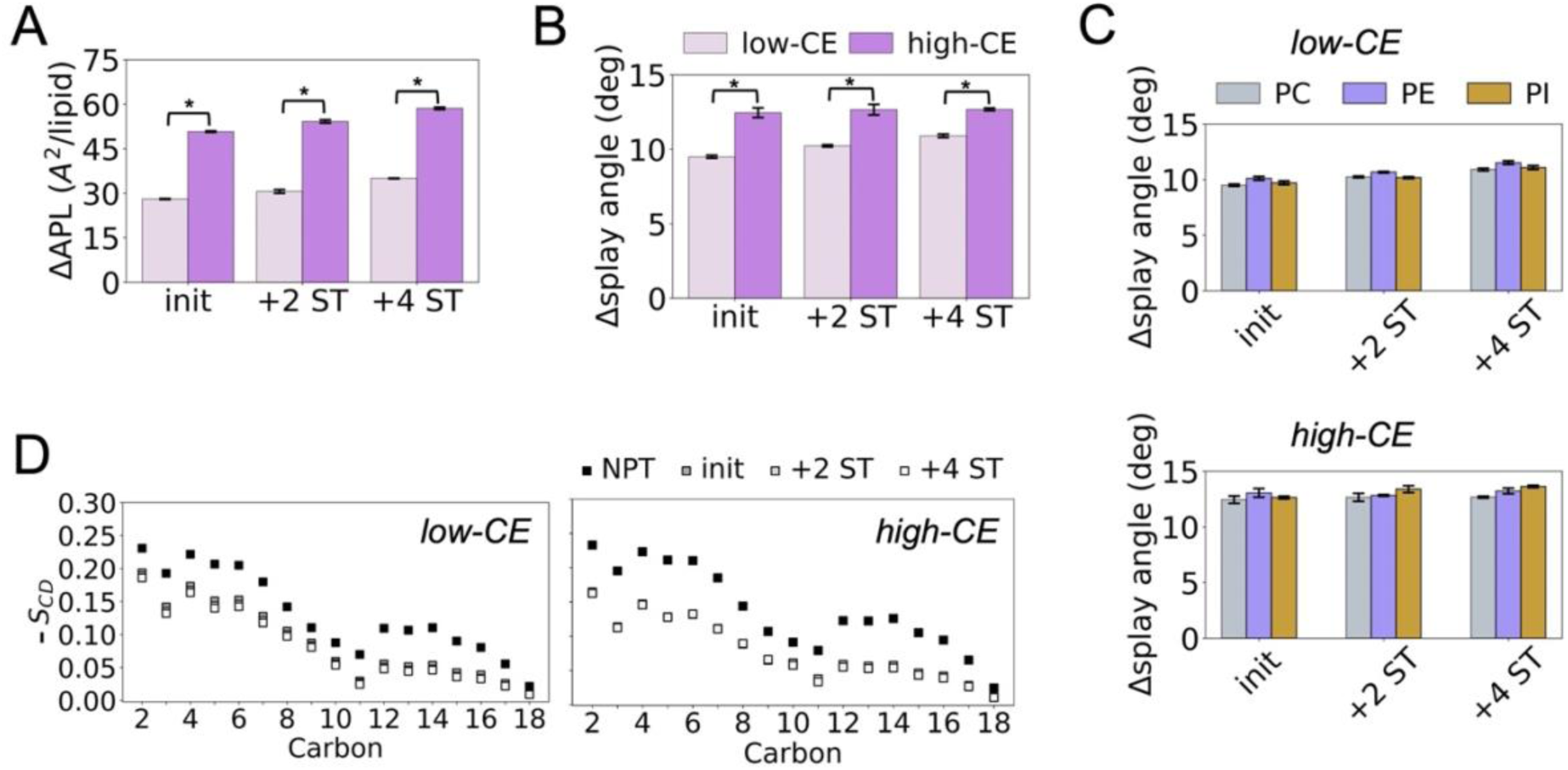
Structure of monolayer lipids in the presence of CE. **A)** Area per lipid. **B)** PC lipid splay angle **C)** Comparison of PC, PE and PI lipid splay angles in *low-CE* and *high-CE* models. **D)** Deuterium order parameters (S_CD_) of PC lipid sn1 tails in *low-CE* and *high-CE* LD models. Larger -S_CD_ indicates more ordered packing, smaller -S_CD_ indicates more disordered packing. Delta (Δ) indicates the difference calculated between the measured value and the *npt* case of each model. Standard error across replicas indicated with error bars, and “*” indicates significant difference in means (p < 0.05) between compared models (*low-CE* and *high-CE*).

In the presence of CE, stretching induces compression of the aqueous and NL phases, as seen in the EDP profiles of CE and water for the *high-CE* model (Fig. 5A-B). This compression is less pronounced in the *low-CE* model (Fig. S8). Increase in applied tension promotes repulsion of water from the monolayer headgroups, with no notable differences due to lower or higher CE concentration (Fig. 5C). The degree of water permeation is negative for all cases, highlighting that water is very sparsely found at or below the headgroup level of the monolayers. This contrasts with the stretched *Pure* and *ER* models that some amount of water entry into the monolayer hydrophobic core. Though the higher tension applied in the CE-containing models may influence this, the presence of CE in the model enhances the hydrophobic character of the monolayer, which may drive the stronger water repulson in the *low-CE* and *high-CE* models.

**Figure 5.**
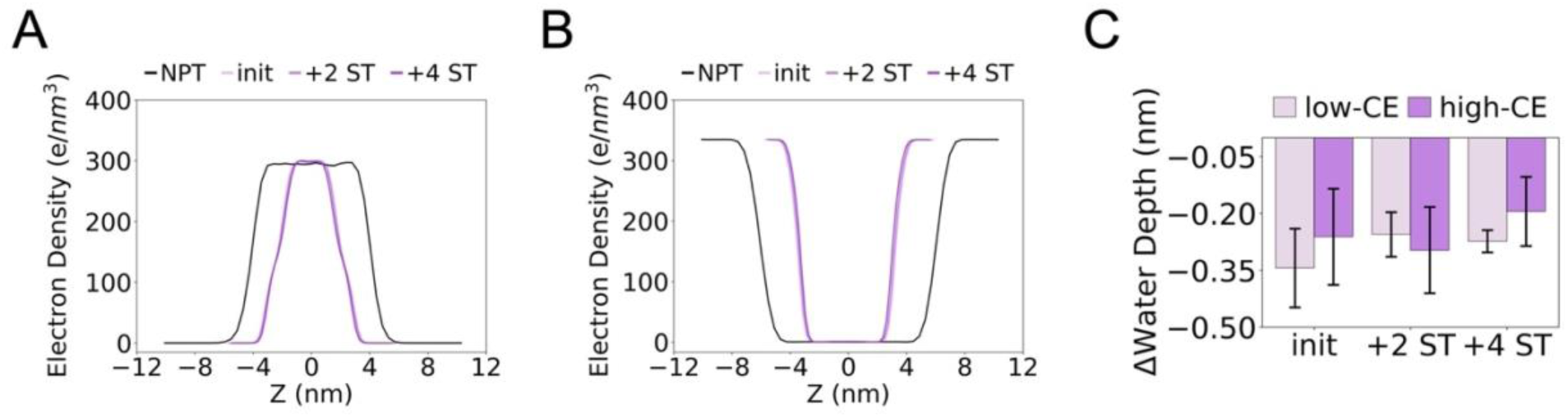
Partitioning of molecular components in the *high-CE* model. **A)** Electron density profile (EDP) illustrates partitioning behavior of CE and **B)** water molecules. **C)** Water depth past phosphate heads in the *low-CE* and *high-CE* models. Standard error across replicas indicated with error bars. No significant difference in means (p < 0.05) between compared models (*low-CE* and *high-CE*).

CE interdigitation has been shown to modulate LD structure across multiple MD simulation works ^24, 25, 27^. In this work, CE interdigitation is defined as when the terminal carbon of either the tail or cholesterol (chol) moiety was found within 10 Å of the monolayer phosphorus atoms (Fig. S9A). The interdigitated number of the bulkier chol-end in the top and bottom leaflets stabilizes at 80 and 50 ns of the *npt* and maximum stretched (*+ 4 S.T.*) runs of the *low-CE* model, and 275 and 250 ns of the *npt* and *+ 4 S.T.* runs of *high-CE* model (Fig. S9B), well within the above noted timescales of equilibration. The number of interdigitated CEs at each applied tension more than doubles after increase in CE concentration (Fig. 6A). In the *low-CE* model, increase in tension promotes both chol-end and tail-end interdigitation, while there is stepwise increase only in the tail-end interdigitation in the *high-CE* model. There is a greater number of interdigitated CEs inserted with the oleate tail compared to the chol-end as CE concentration increases. Prior to stretching, 0.5 and 0.9 mol% of all CE interdigitate in the *low-CE* and *high-CE* models, which is comparable to the 0.3-1.9 mol% penetration of cholesteryl oleate featured in another simulation study ^23^. At maximum stretch conditions, the leaflets contain about 2 and 4 mol% of the total CE within the LD models (Table S3).

**Figure 6.**
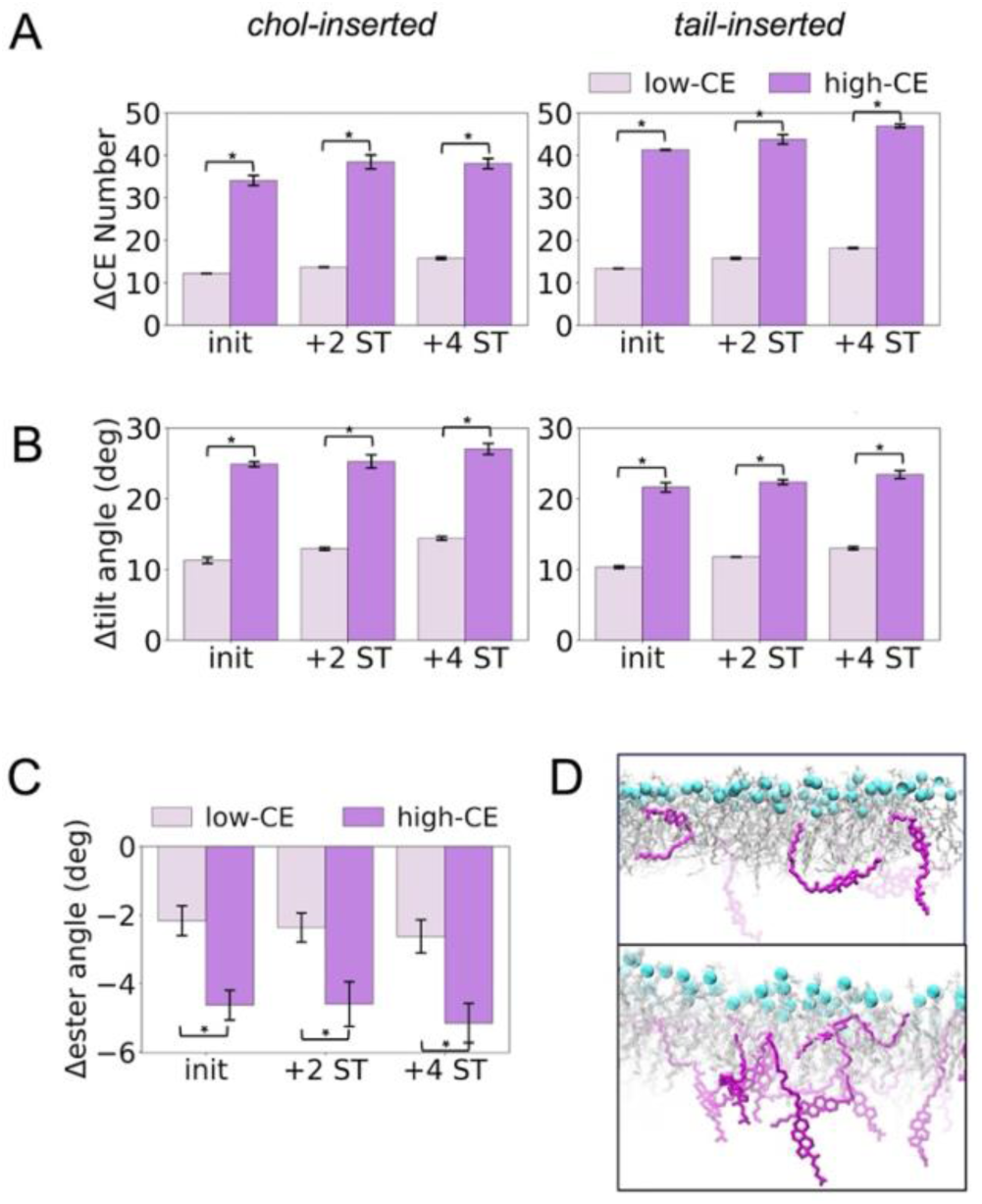
Interdigitated CE behavior in *low-CE* and *high-CE* models. **A)** Number of CEs inserted with the cholesterol and tail ends. **B)** Tilt angle of CEs inserted with the cholesterol and tail ends. **C)** Ester angle of all interdigitated CEs. **D)** Close up snapshots that show orientation of interdigitated CEs in *low-CE* (top panel) and *high-CE* (bottom panel) monolayers. Phospholipid tails colored in grey, P atoms in cyan and CE in purple. Delta (Δ) indicates the difference calculated between the measured value and the *npt* case of each model. *init*, *+2 S.T* and *+4 S.T.* corresponds to the 16, 18 and 20 mN/m cases in the *low-CE* model, and the 28, 30 and 32 mN/m cases in the *high-CE* model. Standard error across replicas indicated with error bars, and “*” indicates significant difference in means (p < 0.05) between compared models (*low-CE* and *high-CE*).

Tilt angle measurements identify the preferred conformations of CEs in terms of the chol moiety or the portion of the oleate tail that begins after the double bond (see Fig. S9C). The tilt angle increases for both chol-end and tail-end interdigitated molecules as CE concentration and applied tension increase (Fig. 6B). However, the tail-end inserted CEs are straighter than the chol-end counterparts with respect to the monolayer normal, suggesting a more ordered degree of packing of the oleate tails regardless of CE abundance. We also computed the ester angle to examine changes in the conformation of the entire CE molecule (Fig. S9D). This angle becomes smaller in both models upon stretching (Fig. 6C). The representative snapshots of interdigitated CE in Figure 6D shows possible conformations in both models, ranging from a near linear and slanted conformation partially inserted into the monolayer, to a folded conformation at the ester angle completely submerged in the monolayer. The monolayer of the *high-CE* model is clearly more saturated with CEs than the *low-CE* model. These trends suggest that CE changes its conformation to fill up vacant spaces in the monolayer, instead of opting for a clean vertical insertion as one may assume based on the crystalline phase formations that have been observed in experiment ^15, 17^. This is in contrast to cholesterol arrangement within bilayers, which aligns vertically to the bilayer normal as its concentration increases enhancing lipid packing and membrane order ^44^.

The tilt and ester angles of interdigitated CEs and representative snapshots show repeatedly that the conformation of these molecules can fold within monolayers. It is possible that this results as an overestimation of the strong attractive forces between the CE oxygen and monolayer lipid headgroup atoms ^45^. Nonetheless, the highest penalty score during estimation of the forcefield parameters using CGenFF ^33^ was 15, which is less than the standard at which further optimization is strongly advised ^46^. Improvement and documentation of steryl ester forcefield libraries will be an important contribution that opens the door to more in-depth studies of their dynamics and function in LDs and other biological lipid structures.

To evaluate CE lipid tail packing in detail, the -S_CD_ of the oleate tail was calculated for both chol-end and tail-end interdigitated CEs. In both the featured *npt* and *+4 S.T.* cases, the -S_CD_ is statistically larger when the oleate tail is inserted compared to when the chol moiety is inserted (Fig. 7), showing that interdigitation leads to more ordered packing of the oleate tail. Also, the *high-CE* model exhibits slightly higher ordered packing compared to *low-CE*, especially in the *npt* case. As tension increases, there is decrease in ordered packing of tail-end inserted CEs versus an increase for the chol-end inserted CEs, particularly for the first 8 carbons starting from the end of the chol moiety (Figs. 7A-B and S10A-B). This shows that the carbons closest to the chol-end also experience an ordering effect when inserted into the monolayer. These results suggest a unique response of CE that depends on which end is interdigitated. The chol-end seems to induce greater packing, similar to cholesterol which has been reported to alter structure by increasing stiffness of bilayers made up of fully saturated lipid tails ^47^. There is more persistent ordered packing induced in the *high-CE* model that contains greater CE concentration, consistent with the near constant tilt and number of interdigitated CEs regardless of higher tensions.

**Figure 7.**
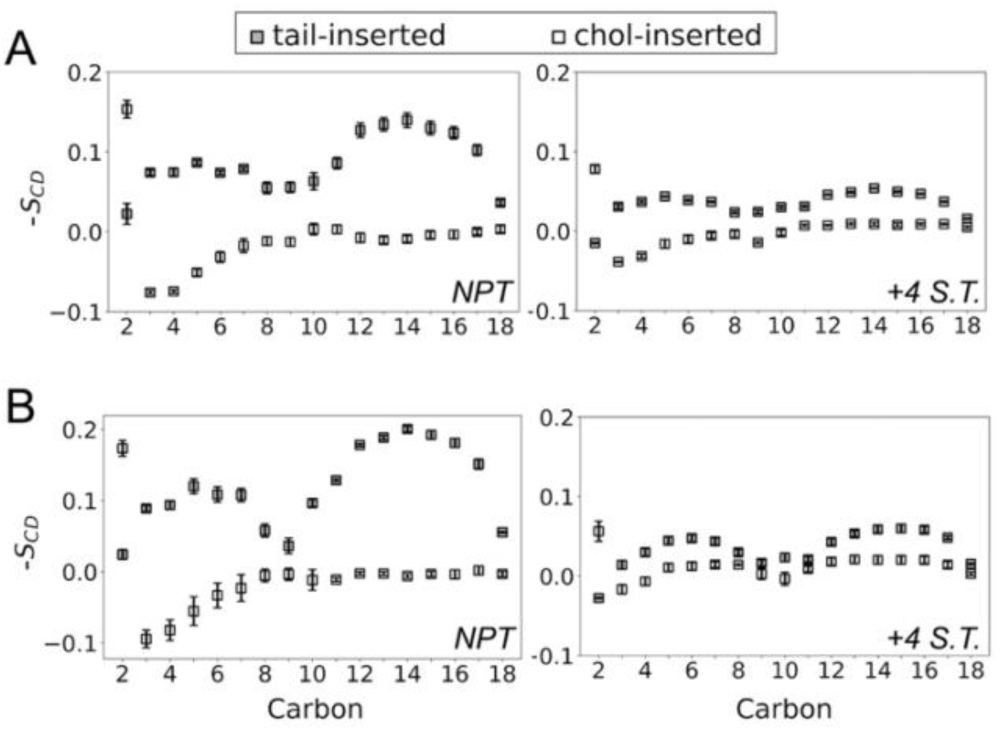
Deuterium order parameters (S_CD_) of CE acyl tail carbons. **A)** Comparison of unstretched (*npt*) and maximum stretched (+4 S.T.) cases in the *low-CE* and **B)** *high-CE* models. *+4 S.T.* corresponds to the *low-CE* 20 mN/m and *high-CE* 32 mN/m cases respectively. Positive, larger - S_CD_ indicates more ordered packing, smaller or negative -S_CD_ indicates more disordered packing. Standard error across replicas indicated with error bars.

The competitive effect of CE abundance and stretching on the monolayer structure is noticeable in all our systems, as each promotes or reduces lipid packing, respectively. Cellular processes such as lipid droplet budding involves an increase in surface tension ^48^. An in vitro study noted the ER lipid composition is an important modulator of surface tension and LD size post-egress, with saturated and mixed PC/PI lipid content fostering the formation of larger LDs with TAGs and steryl esters ^49, 50^. Hence, it is plausible that varying degrees of both CE and surface tension modulate specific LD surface properties under different physiological and pathological conditions.

### CE concentration alters mechanical properties of LDs

Having established the structure modulation trends, we examined how lipid composition influences rigidity and deformation capacity of LDs. Figure 8 summarizes various mechanical properties for the *ER*, *low-CE* and *high-CE* models. Area compressibility, which provides information on the ease of compression or stretching along the monolayer plane, is lower for the *npt low-CE* case compared to the *ER* bilayer (*BL*) case (Fig. 8A). This suggests that the presence of a NL phase between monolayers can reduce the resistance to stretching that is encountered in bilayers. Interestingly, the *high-CE* model has a much larger compressibility of 1.6 N/m – over 5 times that of *low-CE* (0.3 N/m), clearly showing that larger CE concentration works against stretching more than inter-leaflet contact in bilayers.

**Figure 8.**
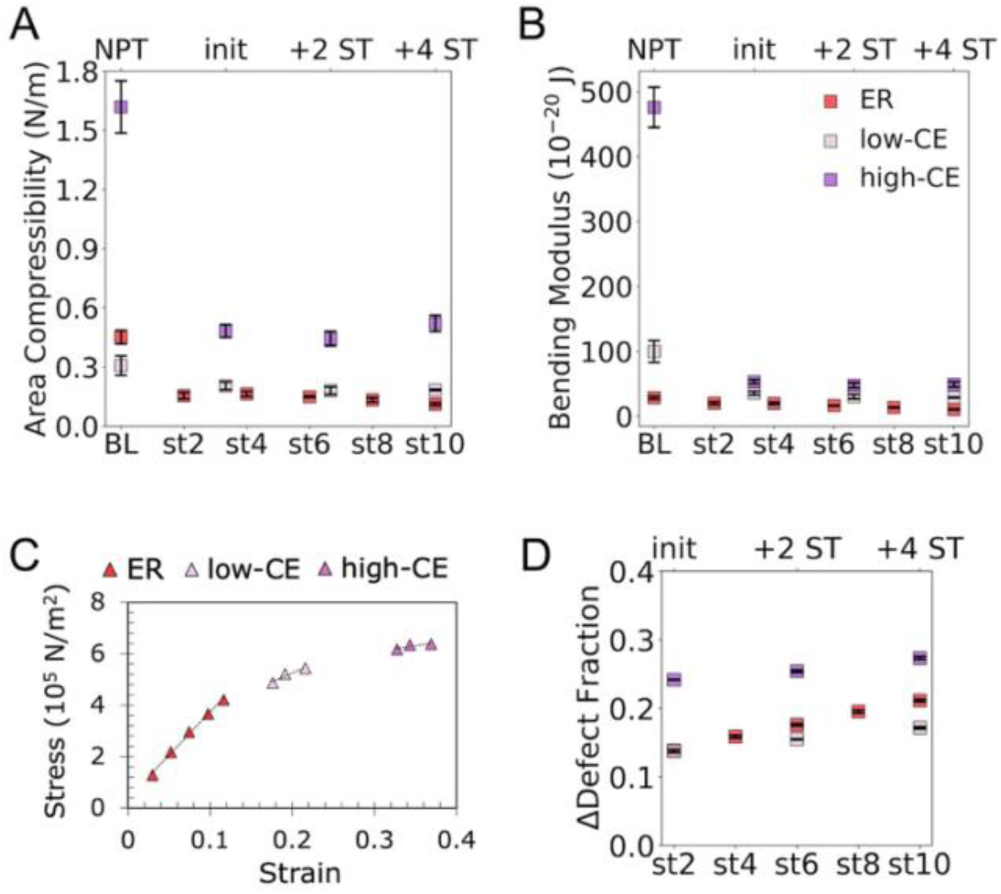
Mechanical and surface properties of modeled LD systems. **A)** Area compressibility, **B)** bending modulus, **C)** stress-strain curve and **D)** defect fraction of monolayer surface in *ER*, *low-CE* and *high-CE* models. Top X axis corresponds to unstretched (*npt*) and stretched cases of the CE-containing models, while bottom X axis corresponds to bilayer (*BL*) and stretched cases in *ER* model. *init*, *+2 S.T* and *+4 S.T.* corresponds to the *low-CE* 16, 18 and 20 mN/m cases, and the *high-CE* 28, 30 and 32 mN/m cases. Delta (Δ) indicates the difference calculated between the measured value and the *BL* (*ER*)/*npt* (*low-CE*, *high-CE*) case of each model. Standard error across replicas indicated with error bars.

The bending modulus is a measure of resistance to change in shape, represented as an energy required to induce a change in curvature ^51^. In this case, both CE-containing models are much more resistant to shape change than the *ER BL* model, with their bending modulus increasing from 28 x 10^-20^ J (*ER npt*) to 100 x 10^-20^ J (*low-CE npt*) and 475 x 10^-20^ J (*high-CE npt)* (Fig. 8B). After stretching, the bending modulus of both CE-containing models reduces sharply, however they remain larger than the stretched *ER* model.

To further quantify the elasticity of LDs, the Young’s Modulus (YM) was estimated via the stress-strain curve shown in Figure 8C. The data shows near-linear increase for the measured stress and strain in the *ER* model. In contrast, there is noticeably lower stress change with applied tension for the *low-CE* and *high-*CE models. YM, estimated from the slope of the curve, decreases by more than a factor of 2 with increasing CE concentration (Table 2). This behavior is similar to a material that approaches its yield strength, which is characterized by a smaller change in the stress compared to the observed elongation – suggesting the LD models become stiffer as CE concentration increases.

**Table 2.** Young’s Modulus measurements in LD models.

| Young's modulus (N/m<sup>2</sup>) |  |
| --- | --- |
| *ER* | $33.518 \times 10^5$ |
| *low-CE* | $13.985 \times 10^5$ |
| *high-CE* | $4.827 \times 10^5$ |

The compressibility and bending resistance of the *high-CE* models are greater than those of the *low-CE* model by at least a factor of 4, which is a clear indication of an increase in rigidity due to CE interdigitation and interaction with the LD monolayer. Mechanical properties of supported lipid bilayers in gel phase exhibit a YM of 2.81 x 10^7^ N/m^2^, area compressibility of 0.2 N/nm and bending modulus of 2.35 x 10^-20^ J ^52^. These bilayers are less stiff compared to the LD models in this work which exhibit ordered arrangement at the monolayer interface prior to stretching (*low-CE, high-CE*: YM = 33.5 x 10^5^, 14.0 x 10^5^ N/m^2^; area compressibility = 0.31, 1.62 N/nm; bending modulus = 100 x 10^-20^, 475 x 10^-20^ J). These results emphasize the influence of CE on inducing tight packing, which markedly increases rigidity of LD monolayers. Therefore, the increase of CE in liver LDs due to cancer likely increases the toughness of the LD exterior, and also influences the interaction with other biomolecules and cellular processes. These changes are likely deleterious to normal breakdown of LDs, and may promote the accumulation of these organelles in liver cancer patients. Further studies are needed to explore this proposed mechanism of CEs modulation of LDs during cancer progression.

The fraction of global lipid packing defects at the surface of the monolayer was examined with the approach outlined by Wildermuth et al ^53^; this involved imaging of the NLs with a different color than lipid headgroups and estimating the exposed NL area as a fraction of total surface area via OpenCV ^54^. Stretching induces a steady increase in the lipid packing defect fraction in all three models (Fig. 8D). However, the defect exposure is greater in the *ER* than in the *low-CE* model, but highest in the *high-CE* model. This non-linear correlation between CE concentration and surface defects may imply a CE concentration threshold in LDs which modulates higher degrees of internal lipid exposure. It is likely that lower CE concentration allows for better fill up of vacancies in the monolayer space by CEs during stretching, but a critical CE concentration saturates the leaflet and promotes disordered crowding at the monolayer-water interface. The alteration of surface defects has known implications on LD protein binding, which occurs by the sensing of packing defects at the interface by protein sub-structures such as amphipathic helices ^24, 55, 56^. Increased defects implies more access of the surface to proteins, but combined with the stiff mechanics of cancerous LDs, the implication of lipid landscapes that are altered in this way still requires further investigation at the cellular level.

### Lipid headgroup re-sorting at the monolayer-water interface

To further understand dynamics at the monolayer interface, lipid-lipid interactions and aggregation analysis were conducted using radial distribution functions (RDFs) of the lipids’ phosphorous atoms. RDF profiles were computed to examine intra-lipid (same lipid species) and inter-lipid (different species) proximity on the monolayer plane. PC lipids of the *Pure* model behave very similarly to those of the *ER* model – the height of the first solvation shell decreases in the stretched cases compared to the *BL* models, and another solvation shell develops around 1 nm, indicating the development of closely-packed PC aggregates (Figs. S11A-B). The positions of the peaks are also effectively the same in both models as applied tension increases.

The PE-PE RDF of the *ER* model exhibits loss of subsequent solvation shells after stretching (Fig. S11B, middle panel), while the PI-PI RDF displays a clear shift of the distribution to the right (Fig. S11B, right panel). However, instead of a decrease like in other cases, the first solvation shell of the PI-PI profile shifts higher as tension increases; signaling an increased likelihood of closely-packed PI headgroups upon stretching.

The position of the first solvation shell of inter-species PC-PE RDFs remains comparable to that of the *BL* model after stretching (Fig. S11C, left panel). The PC-PI distribution shows a unique shift of the solvation shell to the right (Fig. S11C, middle panel), implying less aggregation of PC and PI lipids plausibly in response to the greater population of closely packed PI lipids. Similar to PI-PI results, increase in tension prompts an increase in the height of the first solvation shell of the PE-PI profiles (Fig. S11C, right panel).

When compared to the RDFs of the *Pure* and *ER* models, the headgroups of PI lipids in the CE-containing models aggregate as tension increases, evidenced by the increase in height of their first solvation shell (Fig. 9A). The PC-PC and PE-PE profiles, instead, show a decrease in the height of their first solvation shell with increasing tension (Fig. S12). Interestingly, the reduction of the first and second solvation shells of PC-PE below 1.00 in the *high-CE* model was not observed in any of the other models (Fig. 9B-C, left). This implies that PC-PE headgroup interactions are noticeably disrupted by the presence of CE. PE-PI profiles show the height of the first solvation shell of *init* case of *low-CE* equal to that of *+ 2 S.T.*, while in the *high-CE* model, *init* is lower than both the *+ 2 S.T.* and *+ 4 S.T.* cases (Fig. 9B-C, right). These differing trends at various CE concentration levels point toward a non-uniform modulation of lipid aggregation by CE in monolayers.

**Figure 9.**
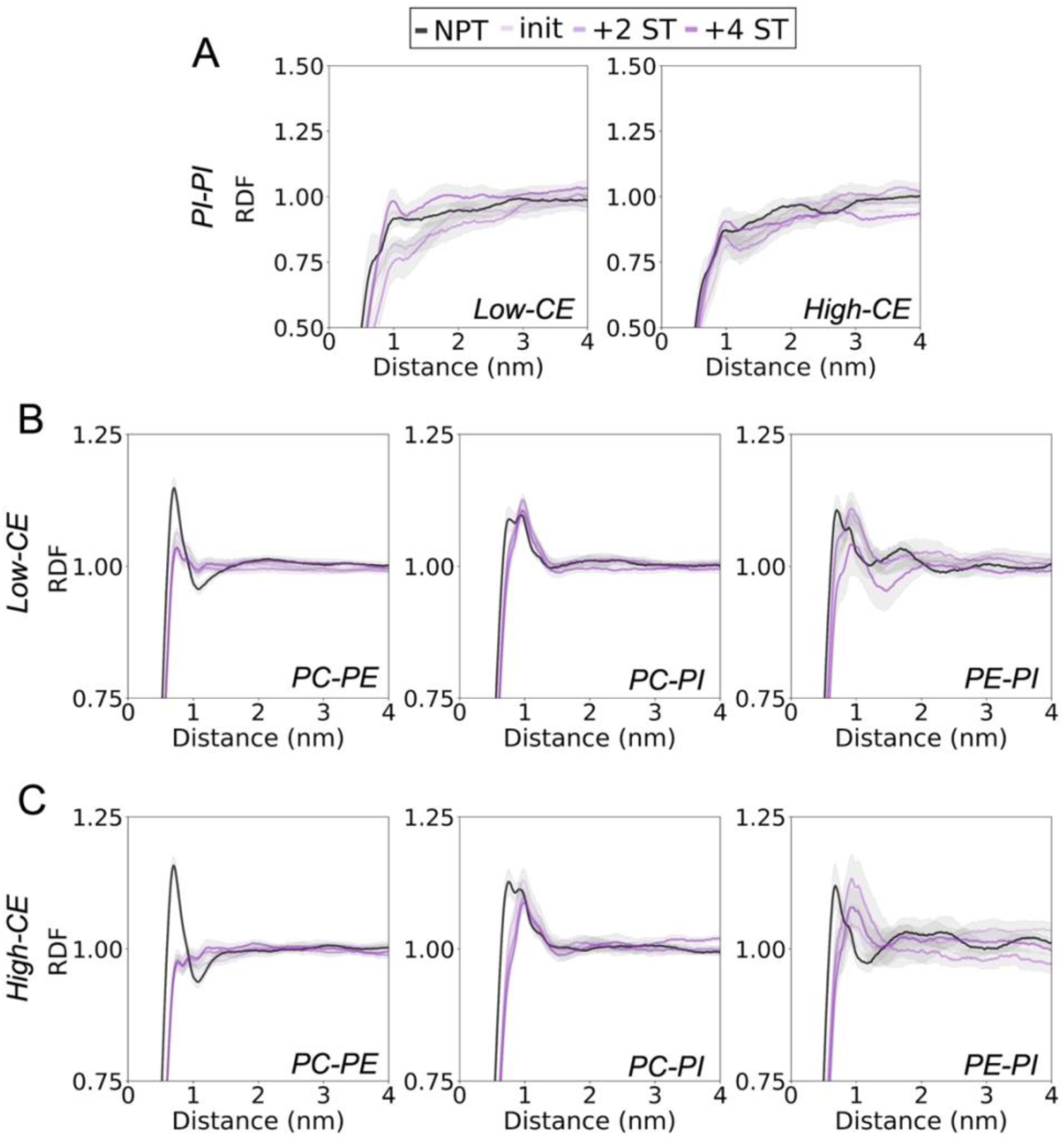
Lipid headgroup 2D RDFs in CE-containing models. **A)** PI-PI RDFs in *low-CE* and *high-CE* models. **B)** PC-PE, PC-PI and PE-PI RDFs in the *low-CE model*. **C)** PC-PE, PC-PI and PE-PI RDFs in the *high-CE* model. Standard error across replicas indicated in grey around individual lines on plots.

Our observations suggest lower headgroup proximity, or loosened packing of lipid headgroups, across most lipid species as stretching becomes pronounced in the absence of CE, except for PE-PI and PI-PI interactions which show increased likelihood for closer packing upon stretching. As CE concentration increases, PC-PE proximity reduces and the strength of PE-PI interactions may become weaker, thereby promoting less resistance to dispersion induced by increased tension. These changes in headgroup packing trends represent an altered landscape of non-covalent interactions at the surface of LDs, which directly influences surface packing defect fraction and subsequent interaction with LD-associated proteins that further regulate LD function.

## Conclusions

There is limited information about CE packing and arrangement at and adjacent to LD phospholipid monolayers, specifically in the event of pathologies that increase CE abundance like liver cancer. Molecular-level understanding is needed on the modulation of structural and mechanical properties of LDs by CE lipids during the progression of liver cancer. Taking inspiration from in vitro micropipette aspiration experiments, the MD simulations conducted in this study innovatively utilized incremental surface tension along the lateral dimensions of trilayer models to investigate the structural, mechanical, and surface properties of LDs at varying concentrations of CEs. Microsecond-long trajectories allowed sufficient sampling to determine monolayers properties in detail, and computational cost was reduced by increasing sampling speed by using hydrophobic DCLE solvent to model the LD core.

From the results, it is seen that CEs interdigitate and alter the packing, water hydration, and mechanical properties of monolayers in unique ways. Increase in CE concentration favors monolayer entry of the acyl tails of CEs over the chol-end to fill vacant spaces within the leaflets, even if chol-end entry promotes more ordered packing as observed in bilayers. Analysis of lipid tail ordering and pairwise aggregation implies that CE interdigitation pushes phospholipid tails and headgroups closer to one another, except for PC/PE and PE/PI headgroups reorganizing farther apart. Measures of resistance to bending, elongation, and compression show increased stiffness and rigidity of LDs with increased CE concentration. This is likely at play during LD accumulation and may encourage reduced LD breakdown in liver cancer cells.

A non-linear correlation between CE concentration and monolayer surface defects is also observed, suggesting a need for a critical CE concentration to increase exposure of NLs to the aqueous phase and modulate binding of surface proteins linked to LD function in healthy or impaired conditions. Though substantial sampling of molecular conformations at the NL-monolayer interface was achieved, spontaneous formation of a fully crystalline phase was inaccessible within the timescale of the conducted simulations. This study provides molecular insight into the link between mechanical properties of LDs and their reinforced integrity in cancer due to the unique contribution of CEs under different physiological conditions.

## Associated Content

### Supporting Information

The Supporting Information is available free of charge at [link].

Text description of all-atom simulation settings for running trilayer systems and featured analyses. Tables summarizing system content; simulation settings; interdigitated fraction of cholesteryl oleate. Figures describing surface tension effects on simulation stability; deuterium order parameters of phospholipid and CE acyl tails; electron density profile of water, DCLE and CE molecules; lateral pressure profile of trilayers; change in shape of trilayers; diffusion coefficients and area per lipid in monolayers; Interdigitated CE metrics (number, angles, and deuterium order parameters); inter- and intra- radial distribution functions of PC and PE lipids of *Pure*, *ER* and CE-containing models.

### Data Availability Statement

Data will be made available on request.

## Author Contributions

**O.C.**: Funding acquisition, Investigation, Methodology, Data curation, Formal analysis, Software, Validation, Visualization, Writing – original draft preparation. **C.L.**: Conceptualization, Funding acquisition, Project administration, Supervision. **V.M.**: Conceptualization, Funding acquisition, Project administration, Supervision, Resources, Writing – review & editing.

### Notes

The authors declare no competing financial interest.

## Acknowledgements

We thank Lily Xueqi Li, Angela Elizabeth Dean, Yi-Heng Huang, Diego Hernandez-Saavedra, and Sayeepriyadarshini Anakk for experimental work that informed modeling of LD systems and discussion of results. Support was provided by the National Institute of Diabetes and Digestive and Kidney Diseases, R01-DK130317 and the Cancer Center at the University of Illinois, Urbana-Champaign (CCIL). The simulations featured in this work utilized Bridges-2/Ocean at the Pittsburgh Supercomputing Center and Stampede3/Ranch at the Texas Advanced Computing Center through allocation BIO240088 from the Advanced Cyberinfrastructure Coordination Ecosystem: Services & Support (ACCESS) program, which is supported by U.S. National Science Foundation grants #2138259, #2138286, #2138307, #2137603, and #2138296. Simulations were also run on the resources provided by the Center for Computational Research at the University at Buffalo. O.C. was supported by the University at Buffalo Presidential Fellowship, and the National Institute of Health’s Initiative for Maximizing Student Development Training Grant T32 GM144920 awarded to Margarita L. Dubocovich (PI).

