## supplemental information for "Cholesteryl Esters Modulate Lipid Droplet Rigidity and Monolayer Organization during Liver Cancer Progression"

### Simulation Settings

The trilayer models in this study underwent equilibration and production phases that vary slightly. Table S1 summarizes the slight differences in settings, while the text below describes general settings across all phases.

**Table S1.** Simulation settings applied to various phases of modeled systems.

| System | Simulation Phase | PME cutoff and fourier spacing | vdW cutoffs | Pressure coupling | Temperature controller | Temperature coupling |
|---|---|---|---|---|---|---|
| bilayer | equilibration | 1.2 nm; 0.12 nm | 1.0 nm (soft); 1.2 nm (hard) | semi-isotropic; 5.0 ps | Nose-Hoover [1, 2] | 1.0 ps |
| DCLE box | equilibration | 1.2 nm; 0.16 nm | 1.0 nm | isotropic; 2.0 ps | V-rescale [3] | 0.1 ps |
| CE/DCLE box | equilibration | 1.2 nm; 0.16 nm | 1.0 nm (soft); 1.2 nm (hard) | isotropic; 5.0 ps | Nose-Hoover [1, 2] | 1.0 ps |
| trilayer | NPT production | 1.2 nm; 0.12 nm | 1.0 nm (soft); 1.2 nm (hard) | semi-isotropic; 5.0 ps | Nose-Hoover [1, 2] | 1.0 ps |
| trilayer | $NP_z\gamma T$ production | 1.2 nm; 0.12 nm | 1.0 nm (soft); 1.2 nm (hard) | surface-tension; 5.0 ps | Nose-Hoover [1, 2] | 1.0 ps |

All simulations used the CHARMM36m forcefield [4, 5] to model forces and interactions of lipids, water and ions. The structure of water molecules followed the three interaction site model, TIP3 [6]. All simulation phases utilized GROMACS MD software [7] to run molecular dynamics. Simulations utilized a timestep of 2 fs and the LINCS algorithm [8] enabled the constraint of covalent-bonded hydrogen vibrations. The Particle Mesh Edwards [9] algorithm modeled electrostatic interactions, while the Verlet [10] algorithm alongside the Lennard-Jones potential force-switch function represented van der Waals (vdW) interactions. Numerical integration utilized the leap-frog algorithm [11] to integrate equations of motion and generate trajectory data. A set temperature of 293.15K captured room temperature conditions during the system runs. This allowed for direct comparison with results from lipid aspiration and other experiments used to determine mechanical and physical properties of LDs in vitro. An applied pressure of 1 bar using the Parrinello-Rahman [12, 13] controller captured atmospheric pressure conditions in the X, Y and Z dimensions in all runs except the $NP_z\gamma T$ production run, where it is applied only in the Z direction. A compressibility of 4.5e-5 $bar^{-1}$ ensured water dynamics similar to those at 1 atm and 300 K. Only the relaxation run involved generation of initial velocities; this was turned off at the start of subsequent equilibration and production runs. During the trilayer NPT production runs, lipid phosphorus atoms in the *low-CE* and *high-CE* models had their positions restrained with a force constant of 500 kJ $mol^{-1}nm^{-2}$. They also had their dihedrals (i.e. angles formed by the glycerol backbone and the first oxygen attached to the C2 carbon) restrained with a force constant of 500 kJ $mol^{-1}rad^{-2}$ to prevent deformation of monolayer leaflets. The trilayer $NP_z\gamma T$ production runs used reference surface tension of 2, 4, 6, 8 and 10 mN/m in the *Pure* and *ER* models, 16, 18 and 20 mN/m in the *low-CE* model, and 28, 30 and 32 mM/m in the *high-CE* model; these choices of surface tension are explained in the Results and Discussion section of the main text.

**Analyses Description**

*Area per lipid (APL):*

Describes the space that individual lipids occupy on the membrane plane, typically the XY plane of the simulation box. It is calculated as follows:

$$APL = \frac{L_x L_y}{\#\ of\ lipids/leaflet} \tag{1}$$

where $L_x$ and $L_y$ are the X and Y dimensions of the simulation box.

*Surface Tension:*

Describes the energy that arises at an interface due to unfavorable interactions between polar and non-polar phases. It is represented as the energy per unit length of an interface. This value is calculated from simulation using the following equation:

$$\begin{aligned}\gamma(t) &= \frac{1}{n}\int_0^{L_z}\left\{P_{zz}(z,t) - \frac{P_{xx}(z,t) + P_{yy}(z,t)}{2}\right\} dz \\ &= \frac{L_z}{n}\left[P_{zz}(t) - \frac{P_{xx}(t) + P_{yy}(t)}{2}\right]\end{aligned} \tag{2}$$

where $L_z$ is the height of the box, $n$ is the number of surfaces, and $P_{xx}$, $P_{yy}$ and $P_{zz}$ are components of the pressure tensor pointing normal to the X, Y and Z plane of the box.

*Lipid tail splay:*

Defined as the angle formed by the two fatty acid tails of a phospholipid. Each tail is assigned as a vector connecting the C2 carbon in the glycerol backbone to the terminal carbon of the acyl chain. Afterwards, the angle between them is calculated as:

$$\theta_{splay} = \cos^{-1}\left(\frac{\boldsymbol{v_1} \cdot \boldsymbol{v_2}}{|\boldsymbol{v_1}||\boldsymbol{v_2}|}\right) \tag{3}$$

where $\boldsymbol{v_1}$ and $\boldsymbol{v_2}$ correspond to the first and second tail vectors.

*Lipid deuterium order parameters (-$S_{CD}$):*

Often used as a measure of ordered packing of lipid tails within a lipid structure. This is defined by the angle formed between the membrane normal vector, usually the Z-axis in the simulation box, and every C-H bond present in the acyl tails of the lipid. The equation to find -$S_{CD}$ is given as:

$$-S_{CD} = -\left[\frac{3}{2}\cos\theta - \frac{1}{2}\right] = -\frac{3}{2}\cos\theta + \frac{1}{2} \tag{4}$$

When $-S_{CD}$ is larger (and positive), it represents more ordered packing, while a smaller (or negative) $-S_{CD}$ represents more disordered packing of lipid tails.

*Lipid diffusion coefficient:*

Describes the speed and ease of lateral movement of lipids. Mean square displacement (MSD), according to the Einstein formula, is given as

$$MSD = \langle \frac{1}{N} \sum_{i=1}^{N} |r - r(t_0)|^2 \rangle \quad (5)$$

where N is the number of system particles the MSD is calculated over, and r are the coordinates in the desired dimension.

Linear regression relates mean square displacement of atoms of individual lipids to lipid diffusion. If we account for movement in only 2 dimensions (X and Y direction),

$$D_{coeff} = \frac{slope(MSD, T)}{4} \quad (6)$$

*Water depth:*

Quantifies permeation depth of water molecules past the monolayer lipid phosphate group, based on the difference between the peaks of the total system EDP and the positions where the water EDP reduces to zero (see figure below).

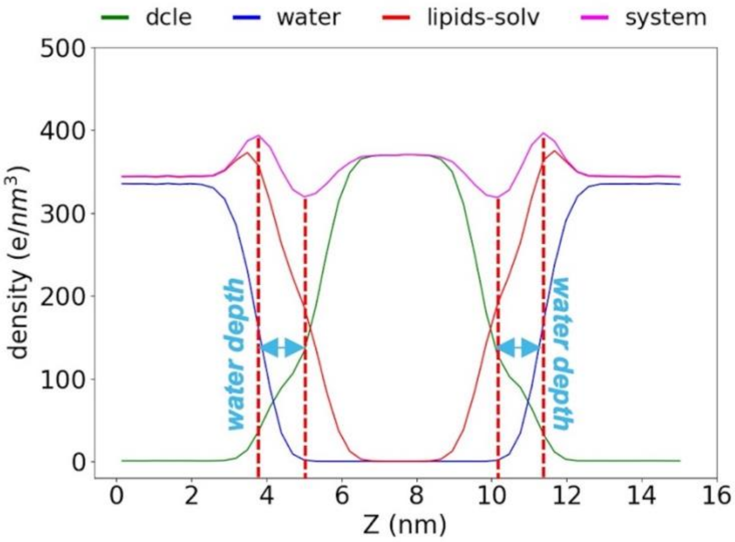


*Area compressibility ($K_{compress}$):*

Describes ease of compression or stretching of monolayers and bilayers. This value is determined using APL data averages and variances in the following equation:

$$K_{compress} = k_B T \frac{APL}{{\sigma_{APL}}^2} \qquad (7)$$

where ${\sigma_{APL}}^2$ is APL variance, $k_B$ is the Boltzmann constant, and $T$ is temperature.

*Bending Modulus ($K_{bend}$):*

Describes energy required to induce curvature in monolayers and bilayers. This value depends on the $K_{compress}$ and can be estimated according to the polymer brush model [14, 15]:

$$K_{bend} = \beta K_{compress} {d_{pp}}^2 \qquad (8)$$

where $\beta = 1/12$ (in bilayer cases) or $1/24$ (in trilayer cases), and $d_{pp}$ is the distance between phosphorus head groups.

*Young's Modulus (YM):*

Describes the elasticity of a material via a stress-strain plot. The equations for stress and strain are given as:

$$Stress = \frac{measured\ surface\ tension}{L_{init}} \qquad (9)$$

$$Strain = \frac{L_{final} - L_{init}}{L_{init}} \qquad (10)$$

where $L_{init}$ and $L_{final}$ are the initial and final horizontal length of the box.

**Table S2**. Summary of simulated LD systems.

| Model | System | Total sim. time (ns) | No. of replicas | Average total atoms | Average box size (after equil. or NPT) (*x, y, z,* in nm) |
|---|---|---|---|---|---|
| ***Pure*** **(400 lip/leaflet)** | DOPC Bilayer (BL) | 200 | 3 | 218,562 | 16.5 x 16.5 x 7.7 |
| | npt | 100 | 3 | 483,821 | 18.5 x 18.5 x 15.9 |
| | 2 mN/m (st2) | 200 | | | |
| | 4 mN/m (st4) | 200 | | | |
| | 6 mN/m (st6) | 200 | | | |
| | 8 mN/m (st8) | 200 | | | |
| | 10 mN/m (st10) | 200 | | | |
| ***ER*** **(400 lip/leaflet)** | PC/PE/PI bilayer (BL) | 200 | 3 | 215,295 | 15.6 x 15.6 x 8.4 |
| | npt | 100 | 3 | 418,689 | 18.1 x 18.1 x 14.7 |
| | 2 mN/m (st2) | 500 | | | |
| | 4 mN/m (st4) | 500 | | | |
| | 6 mN/m (st6) | 500 | | | |
| | 8 mN/m (st8) | 500 | | | |
| | 10 mN/m (st10) | 500 | | | |
| ***ER*** **(600 lip/leaflet)** | PC/PE/PI bilayer | 200 | 3 | 322,945 | 19.4 x 19.4 x 8.3 |
| ***low-CE 60:40 CE:DCLE mass%*** **(600 lip/leaflet)** | npt | 500 | 3 | 861,574 | 20.9 x 20.9 x 20.3 |
| | 16 mN/m (init) | 1,000 | | | |
| | 18 mN/m (+2 S.T.) | 1,000 | | | |
| | 20 mN/m (+4 S.T.) | 1,000 | | | |
| ***high-CE 85:15 CE:DCLE mass%*** **(600 lip/leaflet)** | npt | 1,000 | 3 | 862,850 | 20.1 x 20.1 x 20.7 |
| | 28 mN/m (init) | 2,000 | | | |
| | 30 mN/m (+2 S.T.) | 2,000 | | | |
| | 32 mN/m (+4 S.T.) | 2,000 | | | |

**Table S3**. Total interdigitated fraction of cholesteryl oleate.

| | **Model** | |
|---|---|---|
| **Conditions**$^{+}$ | ***low-CE*** **(mol%)** | ***high-CE*** **(mol%)** |
| **NPT** | 0.51 ± 0.01 | 0.89 ± 0.02 |
| **init** | 1.65 ± 0.01 | 3.77 ± 0.04 |
| **+ 2 S.T.** | 1.82 ± 0.01 | 4.03 ± 0.05 |
| **+ 4 S.T.** | 2.02 ± 0.01 | 4.14 ± 0.03 |

$^{+}$NPT = unstretched conditions.; applied tension cases for low-CE and high-CE respectively: init = 16 and 28 mN/m, +2 S.T. = 18 and 30 mN/m, +4 S.T. = 20 and 32 mN/m.

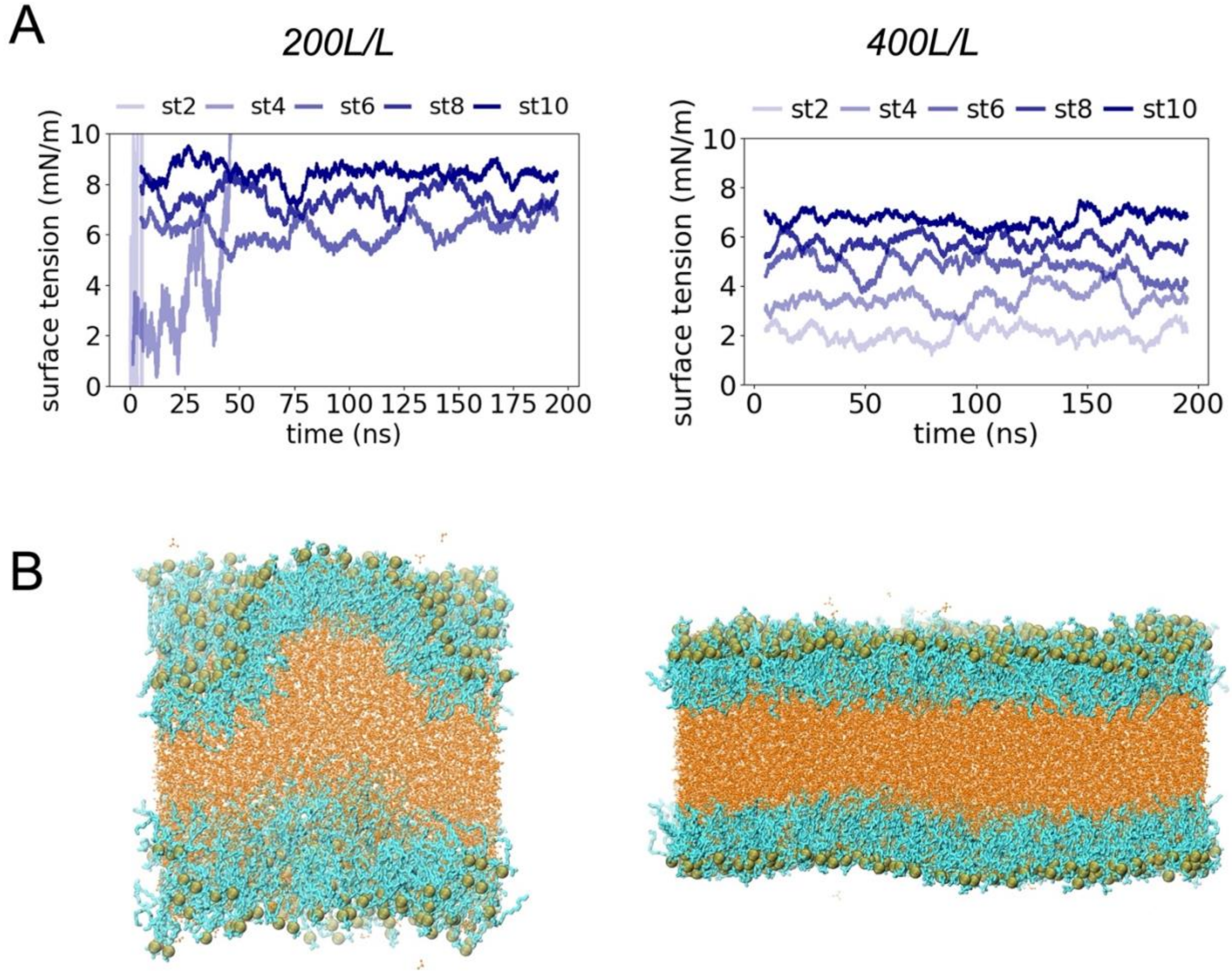


**Figure S1.** Applied surface tension and its effects on simulation stability in the *Pure* model. **A)** Surface tension time series in systems with 200 lipids/leaflet (L/L) and 400 L/L. **B)** Buckling leads to collapse of systems at low tensions in 200 L/L compared to 400L/L. DCLE molecules colored in orange, phospholipids in cyan, and phosphorus atoms in green.

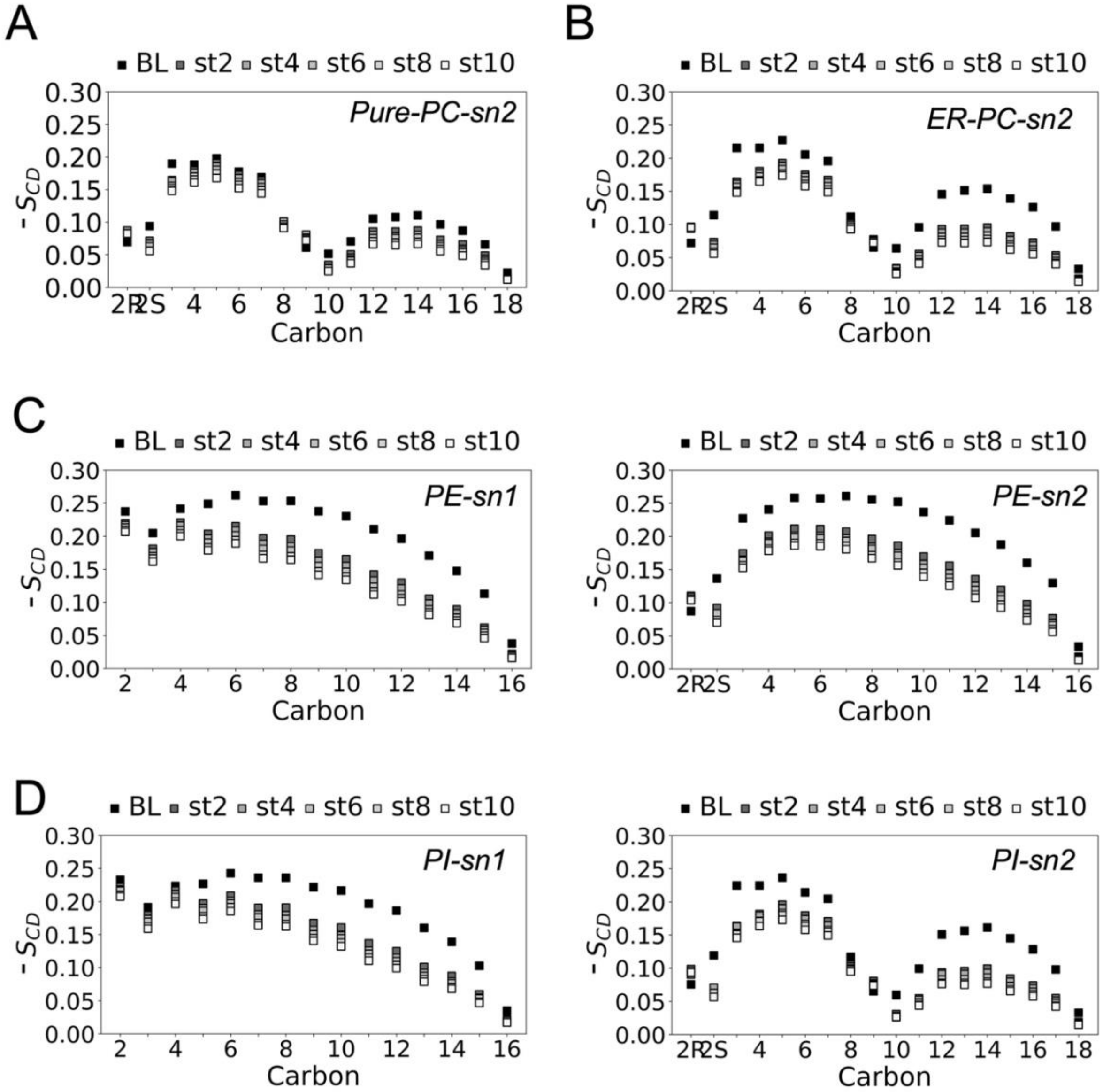


**Figure S2.** Deuterium order parameters ($S_{CD}$) of lipid tails in *Pure* and *ER* LD models. **A)** -$S_{CD}$ of PC sn2 tails in the *Pure* and **B)** *ER* model. **C)** -$S_{CD}$ of PE sn1 and sn2 tails in the *ER* model. **D)** -$S_{CD}$ of PI sn1 and sn2 tails in the *ER* model. Larger -$S_{CD}$ indicates more ordered packing, smaller -$S_{CD}$ indicates more disordered packing.

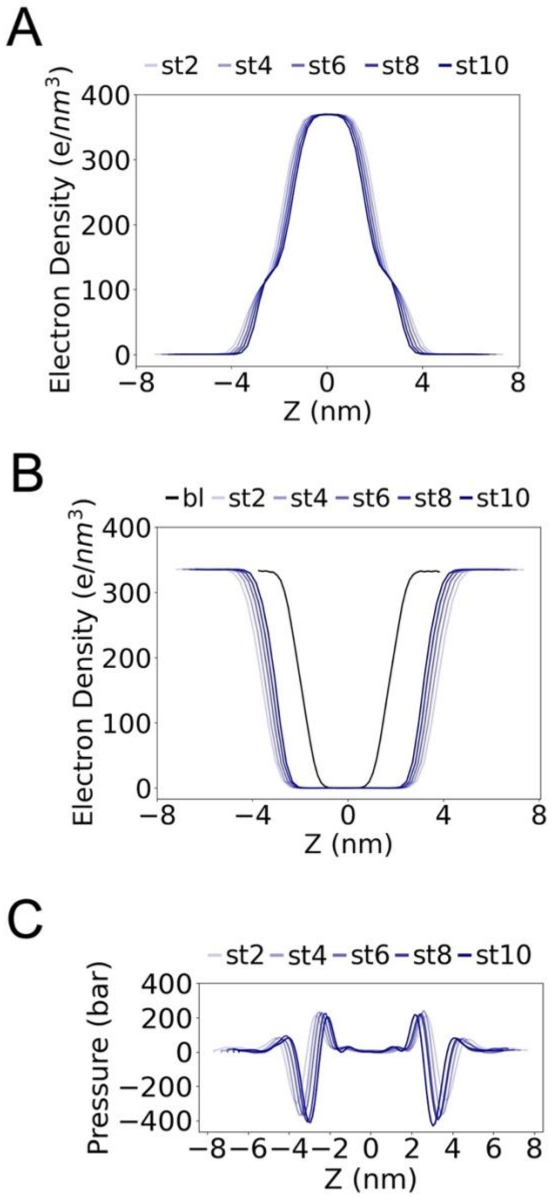


**Figure S3.** Distribution of molecular components in the *Pure* model. **A)** Electron density profile (EDP) of DCLE and **B)** water molecules. **C)** Lateral pressure profile (LPP). Results of increasing tension symbolized by a gradient of light to dark blue.

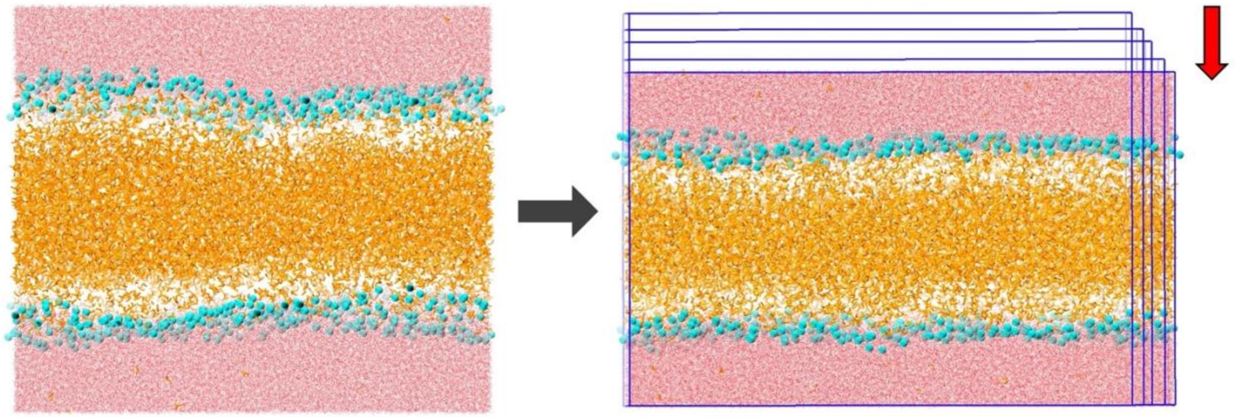

**Figure S4.** Snapshots showing compression (in direction by red arrow) in trilayer systems with increasing surface tension.

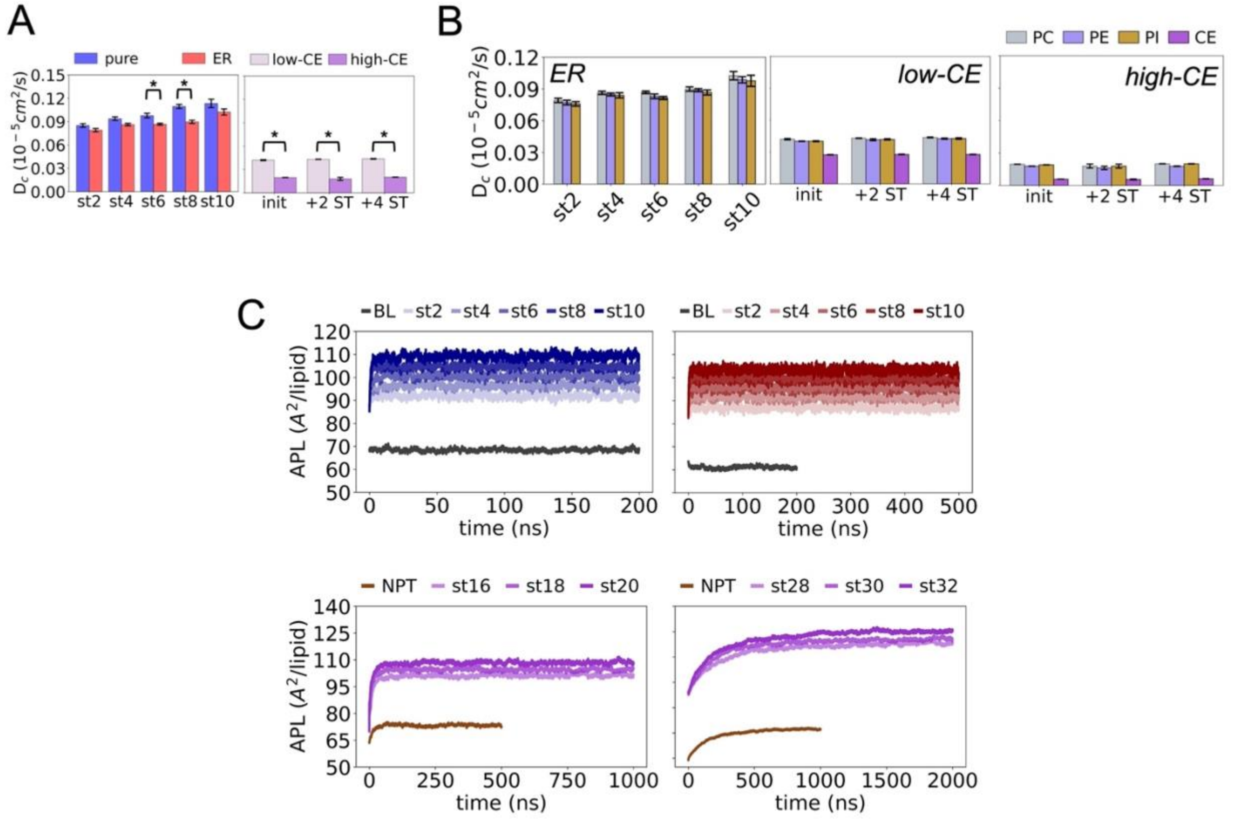


**Figure S5.** Dynamic properties of lipids within simulations. **A)** Diffusion coefficient of PC in all models during stretching. **B)** Diffusion coefficient of all lipid species in stretched *ER* and CE-containing models. **C)** Area per lipid (APL) time series of the first replica of all models. The standard errors across replicas are indicated with error bars, and "*" indicates significant difference in means ($p < 0.05$) between compared models (*pure* and *ER; low-CE* and *high-CE*).

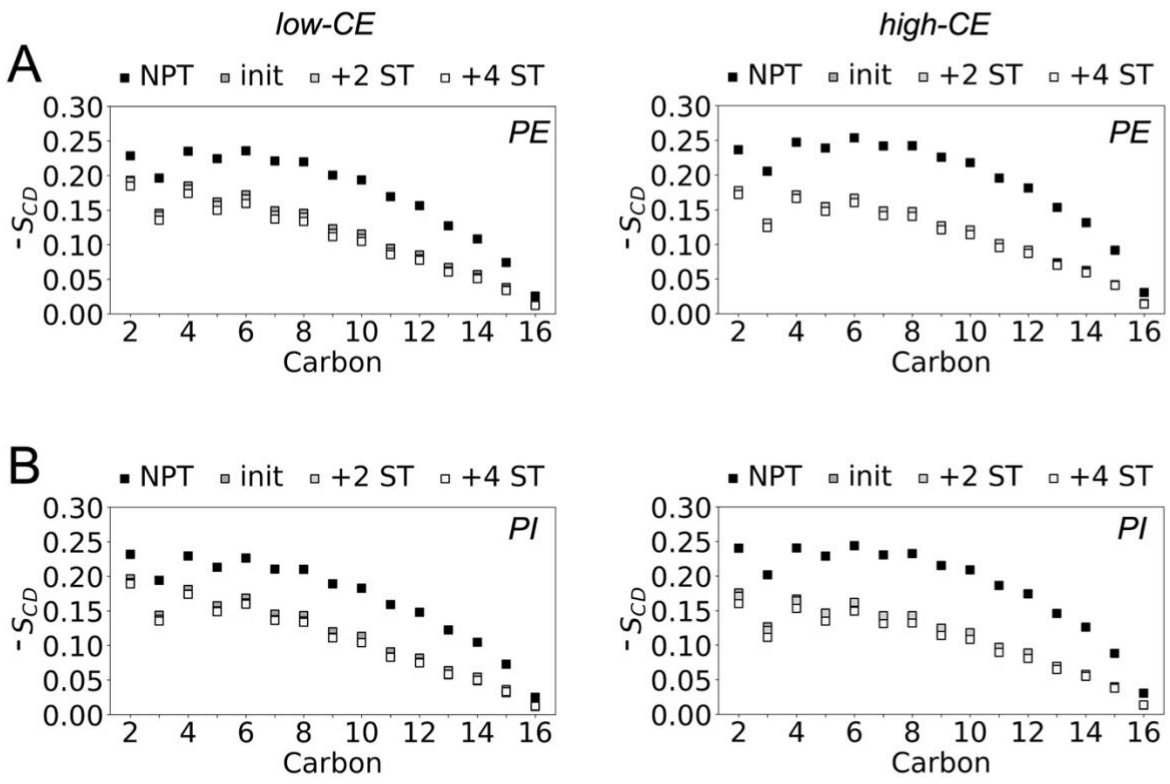


**Figure S6.** Deuterium order parameters ($S_{CD}$) of sn1 lipid tails in *low-CE* and *high-CE* LD models. $S_{CD}$ of **A)** PE, and **B)** PI lipids. Larger -$S_{CD}$ indicates more ordered packing, smaller -$S_{CD}$ indicates more disordered packing.

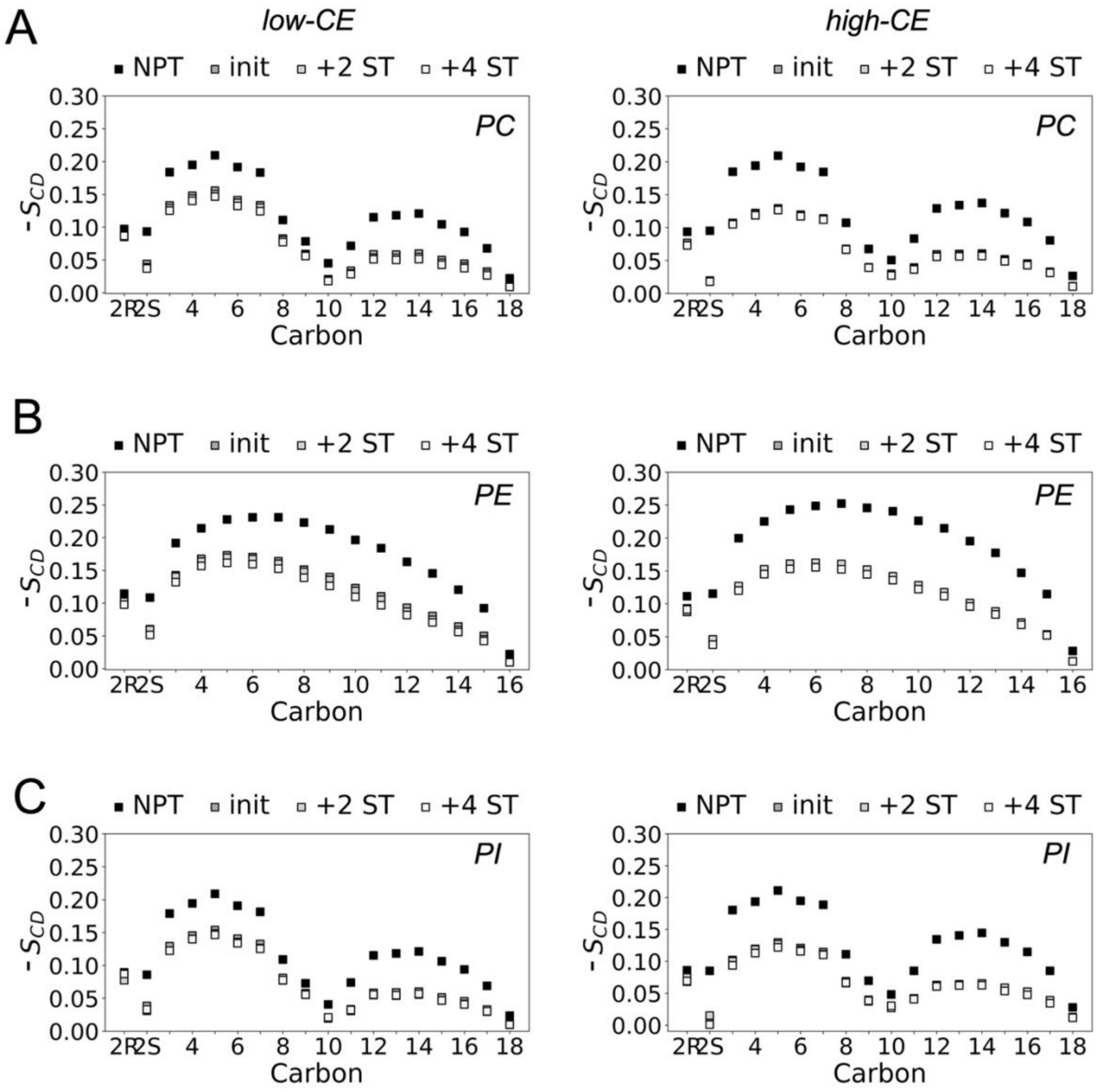


**Figure S7.** Deuterium order parameters ($S_{CD}$) of *sn2* lipid tails in *low-CE* and *high-CE* LD models. $S_{CD}$ of **A)** PC, **B)** PE, and **C)** PI lipids. Larger $-S_{CD}$ indicates more ordered packing, smaller $-S_{CD}$ indicates more disordered packing.

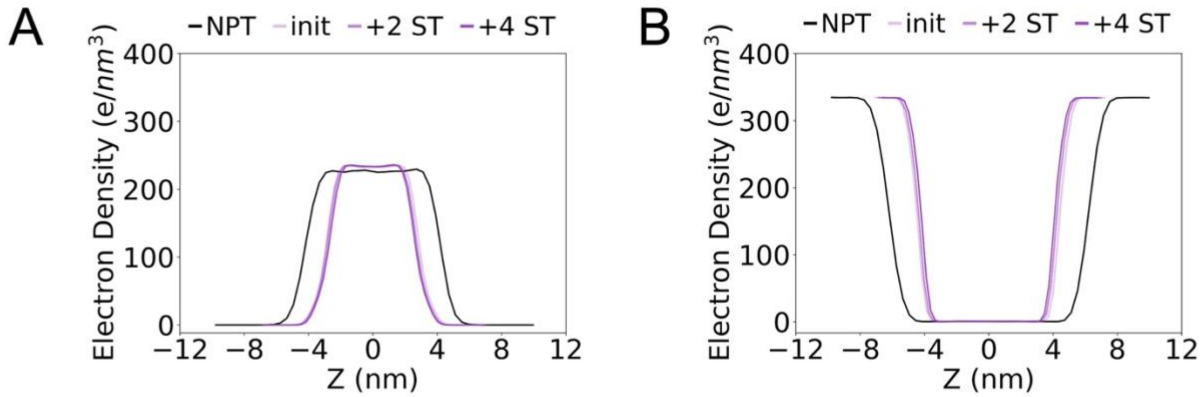


**Figure S8.** Partitioning of molecular components in the *low-CE* model along the z-axis of the simulation box (perpendicular to the monolayer plane). Electron density profiles illustrate partitioning behavior of **A)** CE and **B)** water molecules.

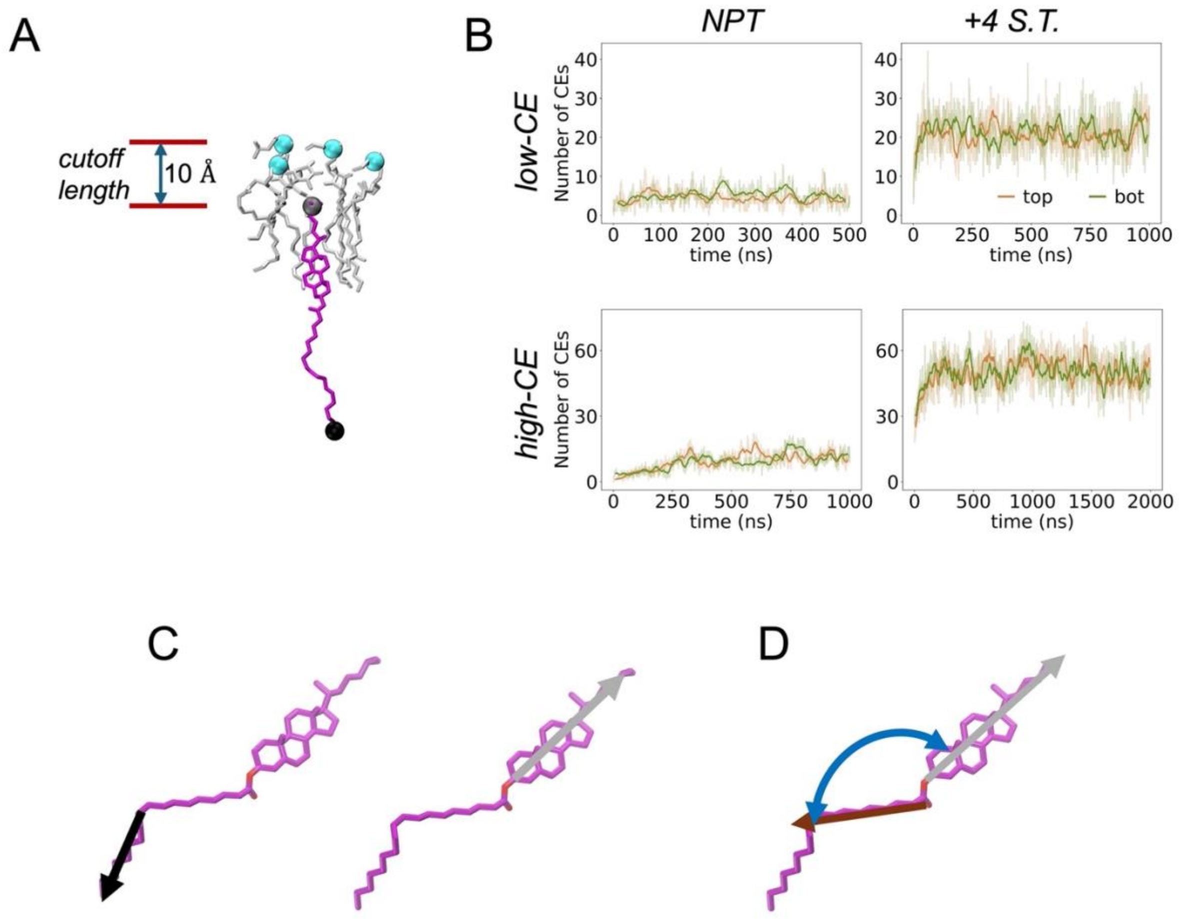


**Figure S9.** Interdigitated CE metrics. **A)** Schematic illustrating interdigitation of CEs (purple; terminal Cs in grey and black) into phospholipid monolayer (silver, Ps in cyan). **B)** Time series of the number of interdigitated CE cholesterol (chol) moieties. Results of *low-CE* and *high-CE npt* and *+4 S.T.* cases indicated in panel. Schematic showing choice of vectors for interdigitated CE angle analysis: **C)** vectors for calculating tilt angle of tail-end and chol-end against the Z normal vector, and **D)** vectors for calculating ester angles. Vector representing acyl tail after double bond indicated in black, the portion before double bond in brown, the chol moiety in grey, and the ester angle in blue.

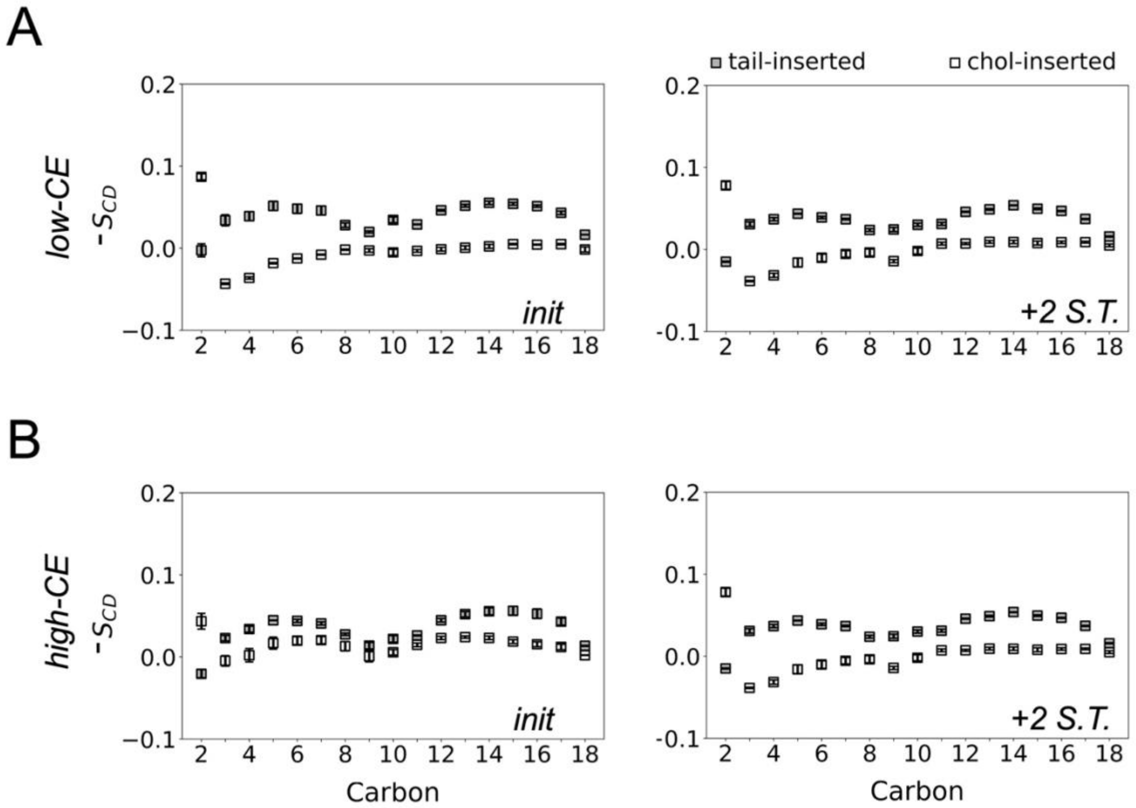


**Figure S10.** Interdigitated CE deuterium order parameters ($S_{CD}$) in **A)** *low-CE* and **B)** *high-CE* models. *init* and *+2 S.T.* correspond to the 16 and 18 mN/m cases in *low-CE*, and 28 and 30 mN/m cases in the *high-CE* models. Positive, larger -$S_{CD}$ indicates more ordered packing, smaller or negative -$S_{CD}$ indicates more disordered packing. Standard errors across replicas are indicated with error bars.

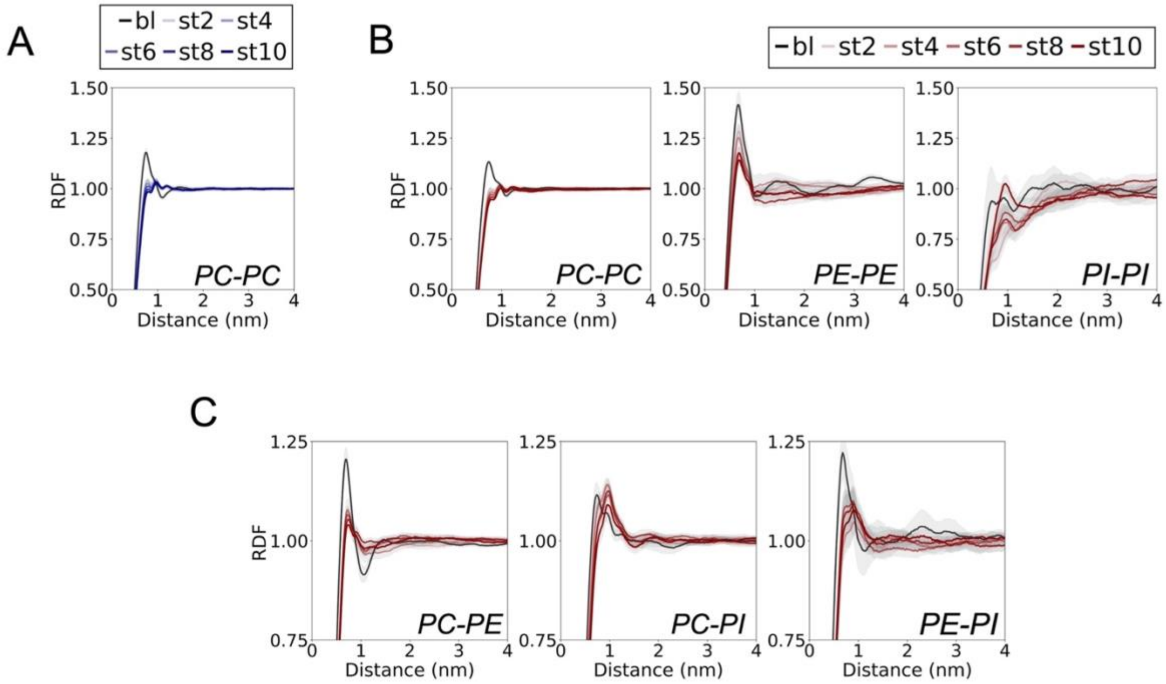


**Figure S11.** Lipid headgroup 2D radial distribution functions (RDFs) in absence of CE. **A)** PC-PC lipid RDF in *Pure* model. **B)** PC-PC, PE-PE and PI-PI RDFs in the *ER* model. **C)** PC-PE, PC-PI and PE-PI RDFs in the *ER* model. Standard error across replicas indicated in grey region around line plots.

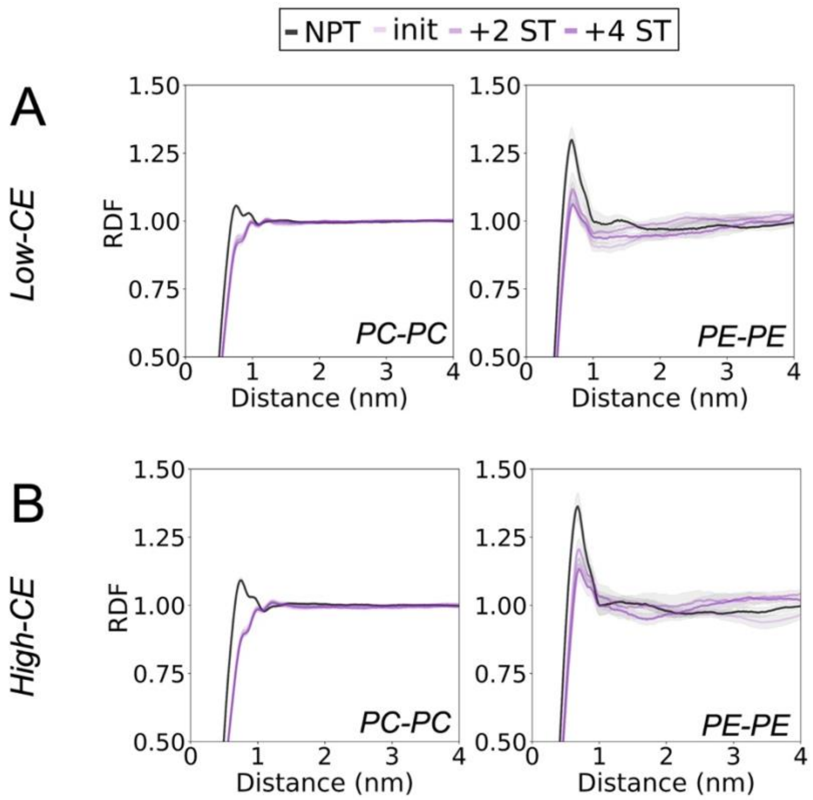


**Figure S12.** Lipid headgroup 2D radial distribution functions (RDFs) in CE-containing models. PC-PC and PE-PE RDFs in **A)** *low-CE* and **B)** *high-CE* models. Standard errors across replicas are indicated in grey region around line plots.
